# A biological model of nonlinear dimensionality reduction

**DOI:** 10.1101/2024.03.13.584757

**Authors:** Kensuke Yoshida, Taro Toyoizumi

## Abstract

Obtaining appropriate low-dimensional representations from high-dimensional sensory inputs in an unsupervised manner is essential for straightforward downstream processing. Although nonlinear dimensionality reduction methods such as t-distributed stochastic neighbor embedding (t-SNE) have been developed, their implementation in simple biological circuits remains unclear. Here, we develop a biologically plausible dimensionality reduction algorithm compatible with t-SNE, which utilizes a simple three-layer feedforward network mimicking the Drosophila olfactory circuit. The proposed learning rule, described as three-factor Hebbian plasticity, is effective for datasets such as entangled rings and MNIST, comparable to t-SNE. We further show that the algorithm could be working in olfactory circuits in Drosophila by analyzing the multiple experimental data in previous studies. We finally suggest that the algorithm is also beneficial for association learning between inputs and rewards, allowing the generalization of these associations to other inputs not yet associated with rewards.

## Introduction

Our brain efficiently commands behavioral outputs based on complex sensory inputs. This process requires transformation of high-dimensional sensory inputs into concise neuronal representations useful for downstream processing. Recent advances in large-scale neuronal recording have highlighted the significance of neuronal population dynamics beyond single neuron activity [1, 2, 3, 4]. Numerous studies have reported that these population dynamics during tasks often operate within a low-dimensional manifold across various brain regions [1, 4, 5]. Low-dimensional representations are expected to enhance robustness against variations and improve generalization [6].

In addition, it has been considered that representations of different categories might be gradually untangled through information processing in the brain [7, 8]. A simple metric for assessing these untangled representations is linear separability, which evaluates the feasibility of distinguishing different objects using a linear hyperplane. Linearly separable representations enable a straightforward downstream readout, while the entangled ones are more difficult to distinguish. Thus, originally entangled high-dimensional inputs are likely transformed into untangled low-dimensional representations [5, 9].

Such representations have also been studied in the context of artificial neural networks. Deep neural networks are considered to solve complex tasks by acquiring useful untangled representations in a supervised manner [5, 10].

The dimensionality of learned representations in deep neural networks was reported to be expanded in the initial layers close to inputs and reduced in the latter layers close to outputs [11]. These findings suggest that deep neural networks acquire untangled low-dimensional representations of input patterns to solve complex tasks. However, supervised learning usually requires a large number of labels, while the brain can learn more efficiently with fewer (or sometimes without) labels. The necessity of backpropagation for training deep neural networks also challenges their feasibility in biological systems, although some recent studies proposed alternative methods [12, 13]. The potential performance of simpler biological circuits than deep neural networks for obtaining good representations, especially in an unsupervised manner and without backpropagation, has not been sufficiently addressed.

Previous works have proposed that simple neural network models can implement linear dimensionality reduction methods such as principal component analysis (PCA) by a biologically plausible synaptic plasticity rule [14]. However, a variety of more complex structures, including entanglements of data manifolds, cannot be extracted by such linear dimensionality reduction methods. To capture the manifold structure in a high-dimensional space and find latent low-dimensional structure, several nonlinear methods, such as t-distributed stochastic neighbor embedding (t-SNE) and uniform manifold approximation and projection (UMAP), have been developed [15, 16]. Among these algorithms, the t-SNE is based on the relatively simple idea that the similarity matrices of the high-dimensional input and low-dimensional output representations are aimed to be closer. Despite its simplicity, biologically plausible implementations are not known.

Here, we propose a biologically plausible new algorithm, Hebbian t-SNE, which performs as effectively as t-SNE in a simple network mimicking the olfactory circuit in Drosophila. The algorithm encompasses three key features: repeated presentation of inputs in random order, a global factor for comparing input and output changes over time, and a middle layer in the network composed of a large number of neurons with sparse activities. The repetitive presentation of inputs allows the network to infer the input and output similarities required for the t-SNE. The inferred similarities are then compared by the global factor in the model, which regulates synaptic plasticity. The middle layer neurons enable the output representations to move independently of each other since input data is transformed into neuronal activities approximately orthogonal to each other in a high-dimensional space. This network structure is consistent with the olfactory circuit in Drosophila, where low-dimensional output representations are acquired in mushroom body output neurons (MBONs) [17] through a feedforward network with the input layer of projection neurons (PNs) and the middle layer of a large number of Kenyon cells (KCs) with sparse activities [18]. To prove the ability of the Hebbian t-SNE, we first show that the Hebbian t-SNE works as efficiently as t-SNE for data in which linear transformations do not work well, such as entangled rings and MNIST. We then suggest that the Hebbian t-SNE might work in real biological circuits by applying it to experimental data of Drosophila olfactory circuits from previous studies [19, 20]. We finally show that Hebbian t-SNE can conduct reward-association learning similarly to Drosophila, with generalization ability utilizing geometric input structures.

## Result

### Construction of the Hebbian t-SNE

The original t-SNE is based on the minimization of the Kullback-Leibler (KL) divergence between the input and output similarities, as we briefly review below. The t-SNE calculates the input similarity between patterns *X*^*i*^ and *X*^*j*^ as 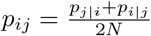, where *N* is the number of total input patterns and *p*_*i*|*j*_ is the similarity of *X*^*i*^ with respect to *X*^*j*^ using the Gaussian measure, which is normalized so that the sum of *p*_*i*|*j*_ over *i* (≠ *j*) is one (Table 1). The output similarity *q*_*ij*_ between output patterns *Y* ^*i*^ and *Y* ^*j*^ is determined based on t-distribution and normalized so that the sum of *q*_*ij*_ over *i* and *j* (*i* ≠ *j*) is one (Table 1). To minimize the KL divergence 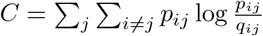 between the input and output similarities, the output representation *Y*^*j*^ are changed according to the negative gradient of the KL divergence 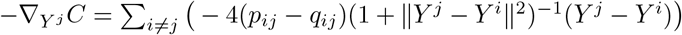 in the t-SNE [15].

**Table 1:**
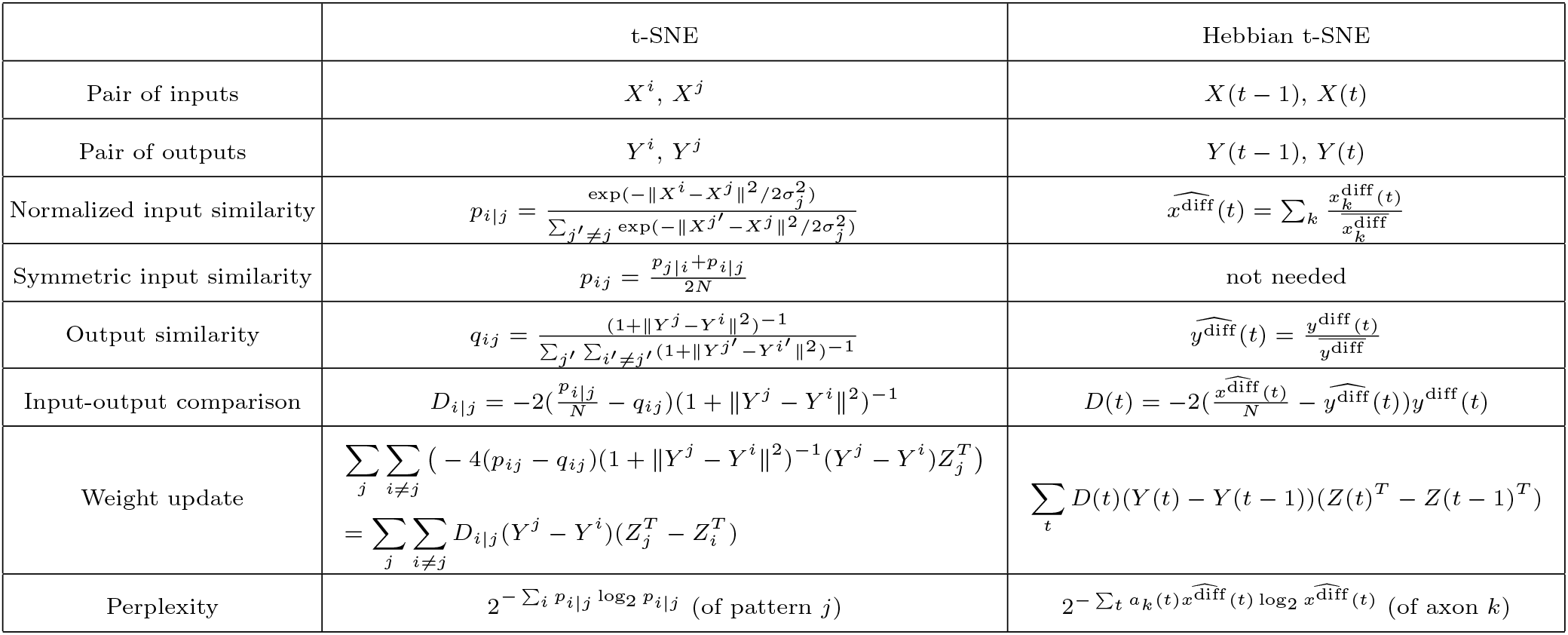
Comparison between t-SNE and Hebbian t-SNE. See Methods for a detailed description of the variables.

To achieve the t-SNE in a biologically plausible way, we considered a simple three-layer feedforward network (Fig. 1a), with the input layer *X*, the middle layer *Z*, and the output layer *Y*. This circuit structure is inspired by the Drosophila olfactory circuit, where *∼* 50 different types of PNs project to *∼* 2000 KCs, which then project to 34 MBONs [18] (Fig. 1b). The input and output layers are comprised of rate neurons equal in their numbers to the input and output dimensions, respectively. In this study, the output dimensions are fixed to two for visualization. The projection from the middle to output layers is linear, *Y* = *WZ*, with plastic synaptic weight matrix *W*. The middle layer contains rate neurons no less than the number of input patterns (or the number of patterns that need to be distinguished; see Discussion) (Fig. 1a). The patterns of middle layer activity *Z* in response to different input patterns are assumed linearly independent. This enables the output representations to have the necessary flexibility for accurate t-SNE; without this linear independence, the output representations are restricted in a subspace with dimensions smaller than the number of input patterns, compromising the accuracy of the t-SNE. To achieve this constraint, we computed the middle layer activity *Z* by projecting inputs *X* by fixed random weights to a high-dimensional space and, then, applied winner-take-all (but see an extension with *k*-winner-take-all in Figs. 4-5 and Extended Data Figs. 4-6, concordant with sparse KC activities in Drosophila [18]). This setting is consistent with the Drosophila olfactory circuit, where the synaptic strength from KCs to MBONs are plastic and regulated by dopaminergic neurons (DANs) while the projection from PNs to KCs is reported to be random and fixed [18, 21] (Fig. 1b).

**Figure 1.**
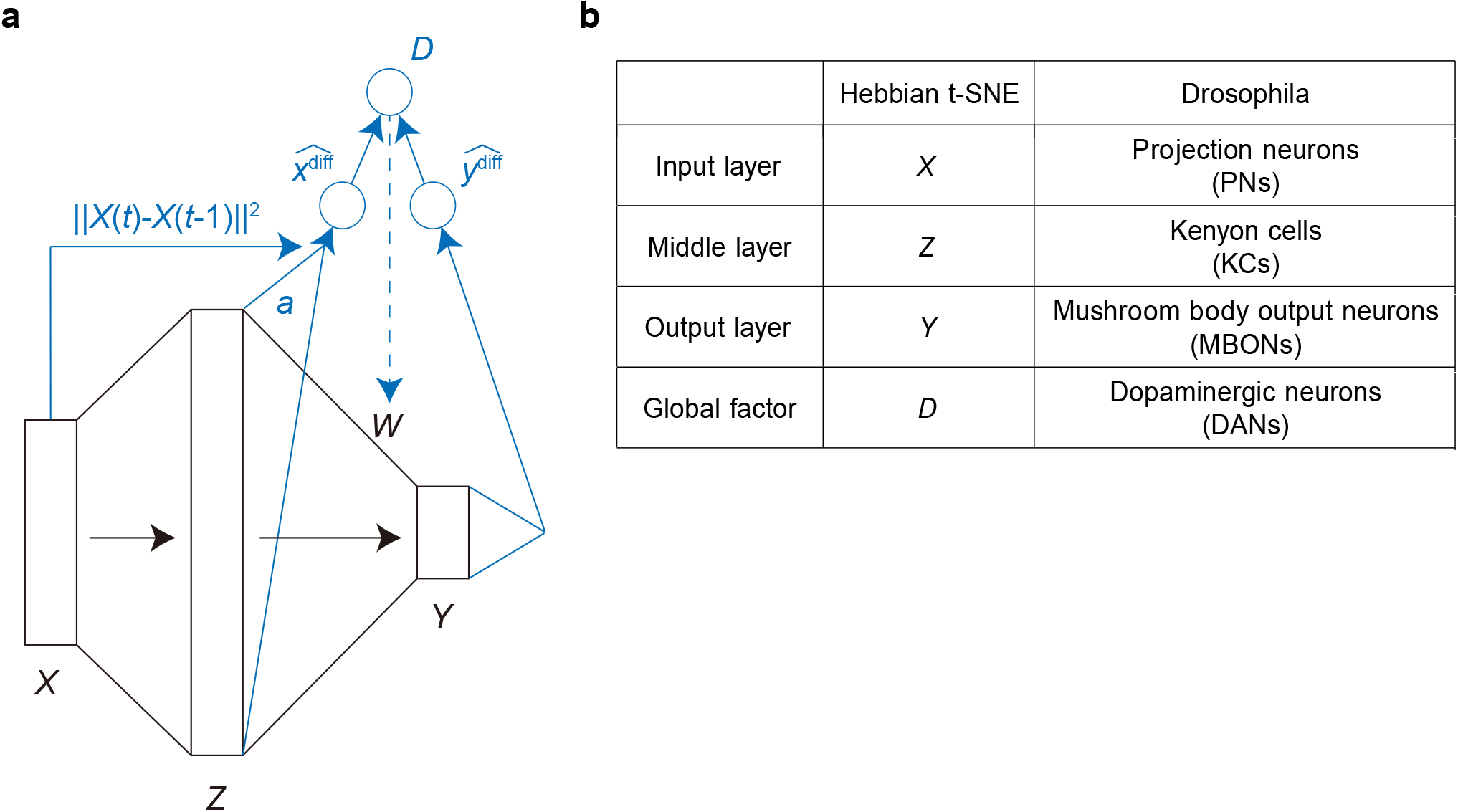
Model structure of the Hebbian t-SNE. (a) Three-layer feedforward model including the input layer *X*, the middle layer *Z*, and the output layer *Y*. The transformation from *X* to *Z* is fixed. The synaptic weight matrix *W* from the middle to output layers is plastic, regulated by the presynaptic and postsynaptic neurons and global factor *D*. The global factor *D* is determined by 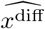 and 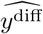 which calculate input and output similarities, respectively. The input similarity 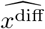 is based on the input difference ∥*X*(*t*) − *X*(*t* − 1) ∥^2^ and axonal signal *a* from the middle layer. (b) Hypothetical elements implementing Hebbian t-SNE in the Drosophila olfactory circuit.

We considered a learning rule of synaptic weight *W* that mimics the t-SNE. We constructed the calculation of the input and output similarities in a biologically plausible way. A detailed mathematical derivation is presented in Methods. We assumed that the whole *N* input patterns *X*^1^, *X*^2^, …, *X*^*N*^ are provided in the random order a large number of times by repeated exposure to the same set of stimuli (or, in some systems, by the neuronal reactivation during sleep [22, 23, 24]). The inputs are provided at each discrete time, denoted by vector *X*(*t*) = (*x*_1_(*t*), *x*_2_(*t*), …, *x*_*r*_(*t*)) for time *t* = 0, 1, 2, …, which elicits the activities of the middle and output activity vectors *Z*(*t*) = (*z*_1_(*t*), *z*_2_(*t*), …, *z*_*s*_(*t*)) and *Y* (*t*) = (*y*_1_(*t*), *y*_2_(*t*)), respectively. In the model, the normalized input similarity *p*_*i*|*j*_ is estimated by neuronal activity 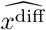, which receives inputs from the middle layer and is modulated by input-difference signal 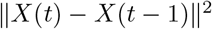 (Fig. 1a). This 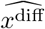 computes how close *X*(*t −* 1) is to *X*(*t*) in the neighborhood of *X*(*t*) according to the input history. Specifically, axonal activity *A*(*t*) = (*a*_1_(*t*), *a*_2_(*t*), …, *a*_*s*_(*t*)) from the middle layer neurons is computed so that *a*_*k*_(*t*) is set to one only at time *t* when the *z*_*k*_(*t*) takes the largest value among the responses to the input patterns *X*^1^, *X*^2^, …, *X*^*N*^ and zero at other times. This axonal output can be biologically computed, for example, with an adaptive firing threshold in each neuron (see Methods). This distinction between the middle layer activity and the axonal activity is utilized to evaluate 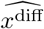 that involves the normalization within the neighborhood of *X*(*t*) (but not of *X*(*t −* 1)). The transmitter release 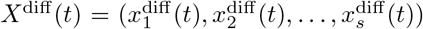 of these axons is modeled as 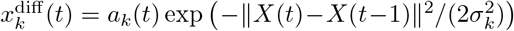 with synapse specific scaling factor *σ*_*k*_. Final<u>ly, th</u>e postsynaptic neuronal activity downstream of these axons is determined to be 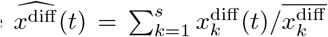 with synaptic weight 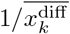. The synaptic weight 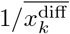 is regulated so that the conditional mean postsynaptic activity given the activation of this synapse (i.e., *a*_*k*_(*t*) *>* 0) is one, similar to the normalization of the t-SNE (Table 1 and Methods). Note that this normalization is straightforwardly computable for each input pattern because the axonal activity *a*_*k*_(*t*) takes a positive value only in response to one input pattern. The output similarity is estimated using *y*^diff^ (*t*) = (1+∥*Y* (*t*)*−Y* (*t−*1)∥^2^)^*−*1^. Then, the term 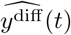, which is obtained by dividing *y*^diff^ (*t*) by its time average 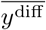, approximates the output similarity *q*_*ij*_ of the t-SNE. The resulting neuronal firing rate *D*(*t*) that integrates the variables 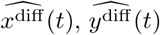 and *y*^diff^ (*t*) is modeled as 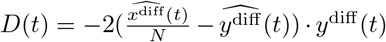, which compares the input and output similarities (Fig. 1a and Table 1). By using this neuronal activity *D*(*t*), the negative gradient of KL divergence between input and output similarities with respect to synaptic weight *w*_*lm*_ is approximated by ∑_*t*_*D*(*t*) *·* (*y*_*l*_(*t*) *− y*_*l*_(*t−* 1)) *·* (*z*_*m*_(*t*) *− z*_*m*_(*t−* 1)). This update rule is biologically plausible. It is given by the product of the presynaptic component *z*_*m*_(*t*)*−z*_*m*_(*t−*1), the postsynaptic component *y*_*l*_(*t*)*−y*_*l*_(*t−*1), and the global factor *D*(*t*). Using this gradient, the synaptic changes are computed by mini-batch and Adam [25]. We call this learning rule Hebbian t-SNE. Note that mini-batch and Adam are considered biologically plausible since they use time-average values of the gradient and its square, which can be computable locally at individual synapses. The model determines the scaling factor *σ*_*k*_ for axon *k* similarly to the t-SNE. The t-SNE selects *σ*_*j*_ of pattern *j* such that the perplexity 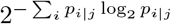 matches the target value determined by the user. The perplexity value, typically set between 5 and 50, determines the number of neighboring points substantially reflected in measuring the input similarity. Mimicking this method, we slowly change *σ*_*k*_ to bring 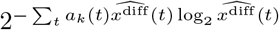 close to the target perplexity. Here, the axonal activity *a*_k_ (*t*) takes a positive value if and only if *X*(*t*) = *X*^*j*^ (see Table 1 and Methods). We let the synaptic weights change after an initial run of 500 steps when the perplexity approached sufficiently close to the target value (Figs. 2e and 3e). We also considered the model in which the calculation of 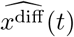 is simplified so that it receives inputs only from the input layer, which we call simplified Hebbian t-SNE (Extended Data Fig. 1). Therefore, we constructed the dimension reduction algorithm imitating t-SNE via a biologically plausible learning rule.

**Figure 2.**
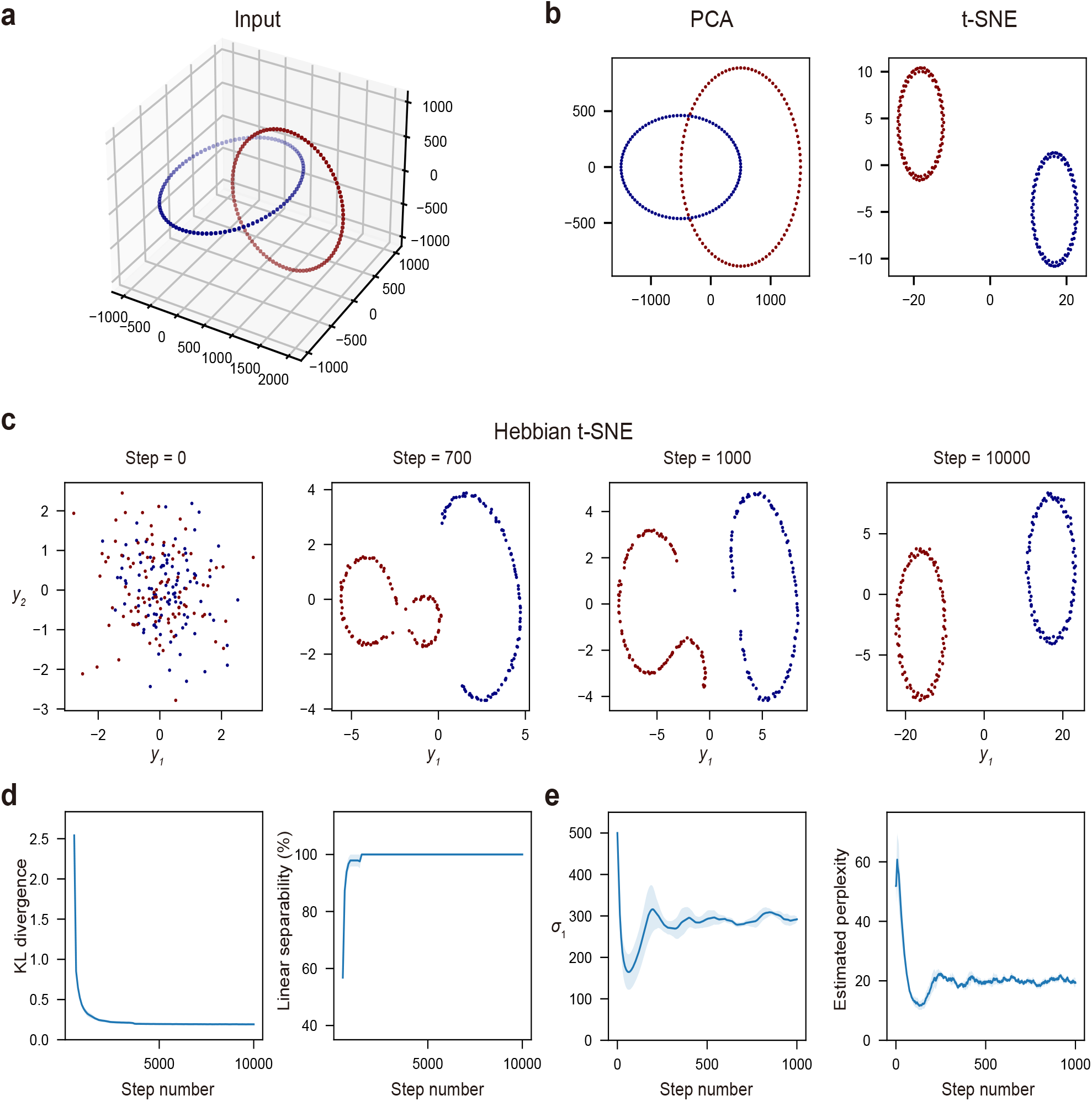
Applying the Hebbian t-SNE to inputs distributed according to entangled rings. (a) Input data. Red and blue dots represent inputs corresponding to each ring. (b) The final two-dimensional output representations obtained by the PCA and t-SNE. While the PCA failed to separate the two rings, the t-SNE successfully separated the two rings. (c) The time course of the representation changes by the Hebbian t-SNE. The two rings were gradually separated along the iterative synaptic weight updates. (d) The changes of the KL divergence between the input and output similarity matrices and the linear separability of two rings in the output representation. The lines and shadows represent the means and SEMs in the five trials, respectively. (e) The changes of the *σ*_1_ value and its related estimated perplexity in the model. The estimated perplexity gradually approached the target perplexity 20 along the changes of *σ*_1_. The lines and shadows represent the means and SEMs in the five trials, respectively.

### Dimensionality reduction of high-dimensional data by Hebbian t-SNE

To demonstrate the effectiveness of the Hebbian t-SNE across various datasets, we first considered the input of two entangled rings (Fig. 2a). While the linear map, such as the PCA, failed to separate entangled rings, the t-SNE successfully provided the separated representation of two rings (Fig. 2b). As with t-SNE, the Hebbian t-SNE succeeded in separating two rings (Fig. 2c). Since the coordinates of the output representations were initialized at random, the KL divergence between the original three-dimensional input and the two-dimensional output similarity matrices took a large value. The linear separability (see Methods for the definition) of two rings in the output representation was small at the initial condition (Fig. 2d). Following the iterative updates of the synaptic weights *W*, the KL divergence between the input and output similarity matrices gradually decreased as expected (Fig. 2d) since the Hebbian t-SNE is constructed to reduce this value. Consequently, the two rings were properly separated and linear separability sufficiently increased (Fig. 2c and d). In this process, the estimated perplexity rapidly approached the set value (20 in this case) in about the first 500 steps as expected (Fig. 2e). Given that the target perplexity determines the spatial scale for measuring input similarity structures, it was set to ensure that input similarity includes information about neighboring points within the same ring while excluding those in another ring. While the appropriate value for target perplexity generally depends on the number of data points and other input data characteristics, we set it between 20 and 40 throughout this manuscript. For the entangled rings, the simplified Hebbian t-SNE also succeeded in separating two rings as well as the Hebbian t-SNE (Extended Data Fig. 2). Therefore, the Hebbian t-SNE can be further simplified in this easy case.

To evaluate the efficacy of the Hebbian t-SNE for more complex higher dimensional data, we considered the MNIST data (Fig. 3a) [26]. As with the entangled rings, the t-SNE but not the PCA provided the low-dimensional representation separating the clusters of different numbers (Fig. 3b). This can be quantitatively assessed using the linear separability, defined as an accuracy rate of solving a multi-class classification problem with the linear support vector machine in a one-vs-one scheme, which was larger in the representation by t-SNE (71 %) than in that by PCA (44 %). Similar to the case of the entangled rings, following the iterative synaptic weight updates, the Hebbian t-SNE provided a low-dimensional representation that distinctly separated the clusters of different numbers (Fig. 3c). More quantitatively, the linear separability rapidly surpassed the PCA level and reached the t-SNE level with learning (Fig. 3d). The estimated perplexity rapidly approached the set value of 40 (Fig. 3e). In contrast, the simplified Hebbian t-SNE generated separated representations only in a limited number of data points, failing to develop good representations (Extended Data Fig. 3). The cluster representing each number was hard to find in the final representation (Extended Data Fig. 3a). Consequently, the linear separability was lower than the Hebbian t-SNE (Extended Data Fig. 3b). Hence, the Hebbian t-SNE, not the simplified t-SNE, obtained a good representation of the MNIST data. Note that although the simplified Hebbian t-SNE was ineffective for complex data such as the MNIST, it might still be adapted in simple biological circuits.

**Figure 3.**
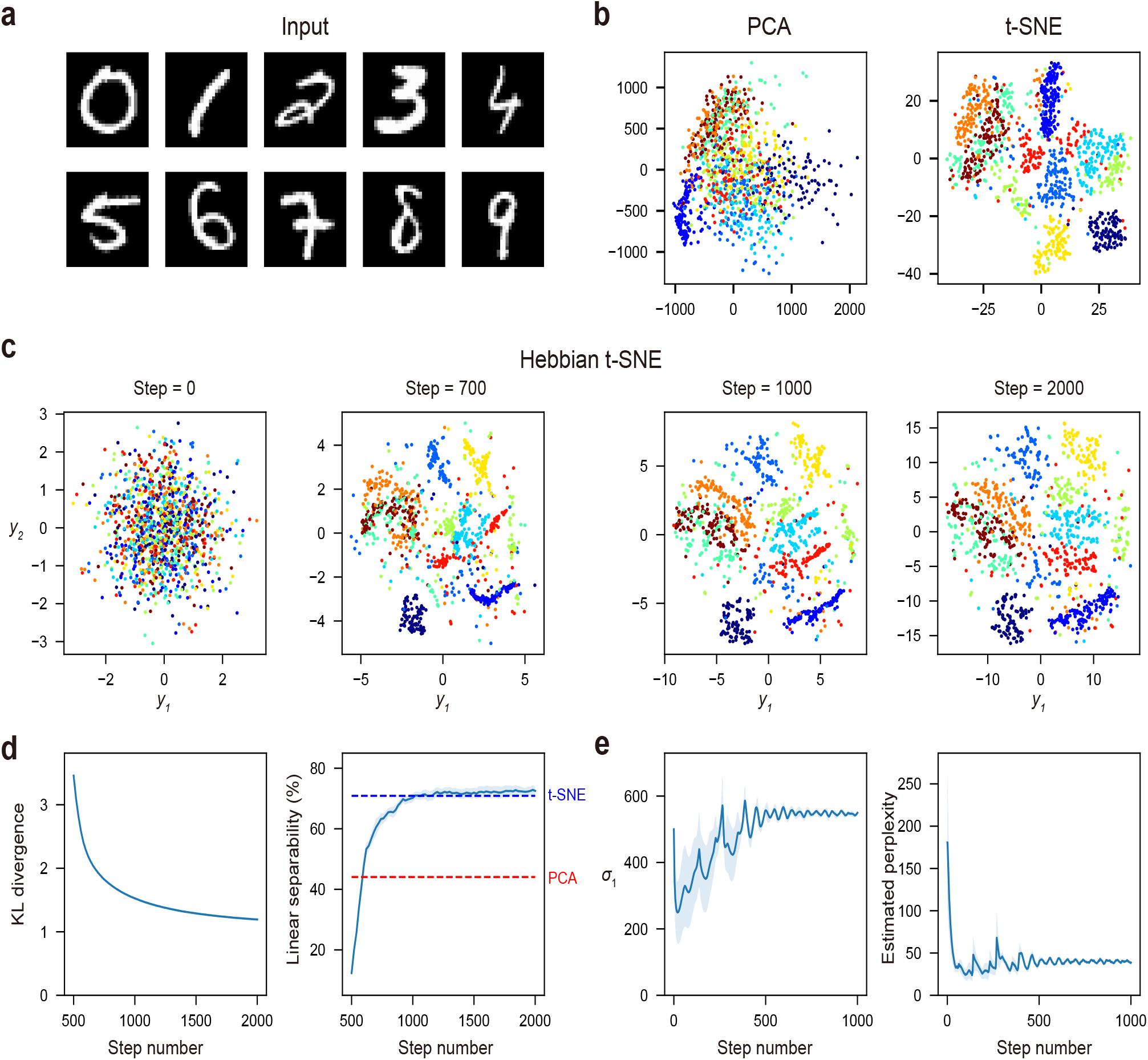
Applying the Hebbian t-SNE to the MNIST data. (a) Sample MNIST data. (b) The two-dimensional representations by the PCA and t-SNE. Each color of the data represents a digit from zero to nine. While the PCA failed to find hidden representations, the t-SNE successfully separated inputs of different digits. (c) The time course of the changes of the representation by the Hebbian t-SNE. The Hebbian t-SNE gradually led to good representations separating different digits. (d) The changes in the KL divergence between the input and output similarity matrices and the linear separability of the ten digits in the output representation. Learning starts at step 500. The lines and shadows represent the means and SEMs in the five trials, respectively. (e) The changes of the *σ*_1_ value and its related estimated perplexity in the model. The estimated perplexity gradually approached the target perplexity 40 along the changes of *σ*_1_. The lines and shadows represent the means and SEMs in the five trials, respectively.

**Figure 4.**
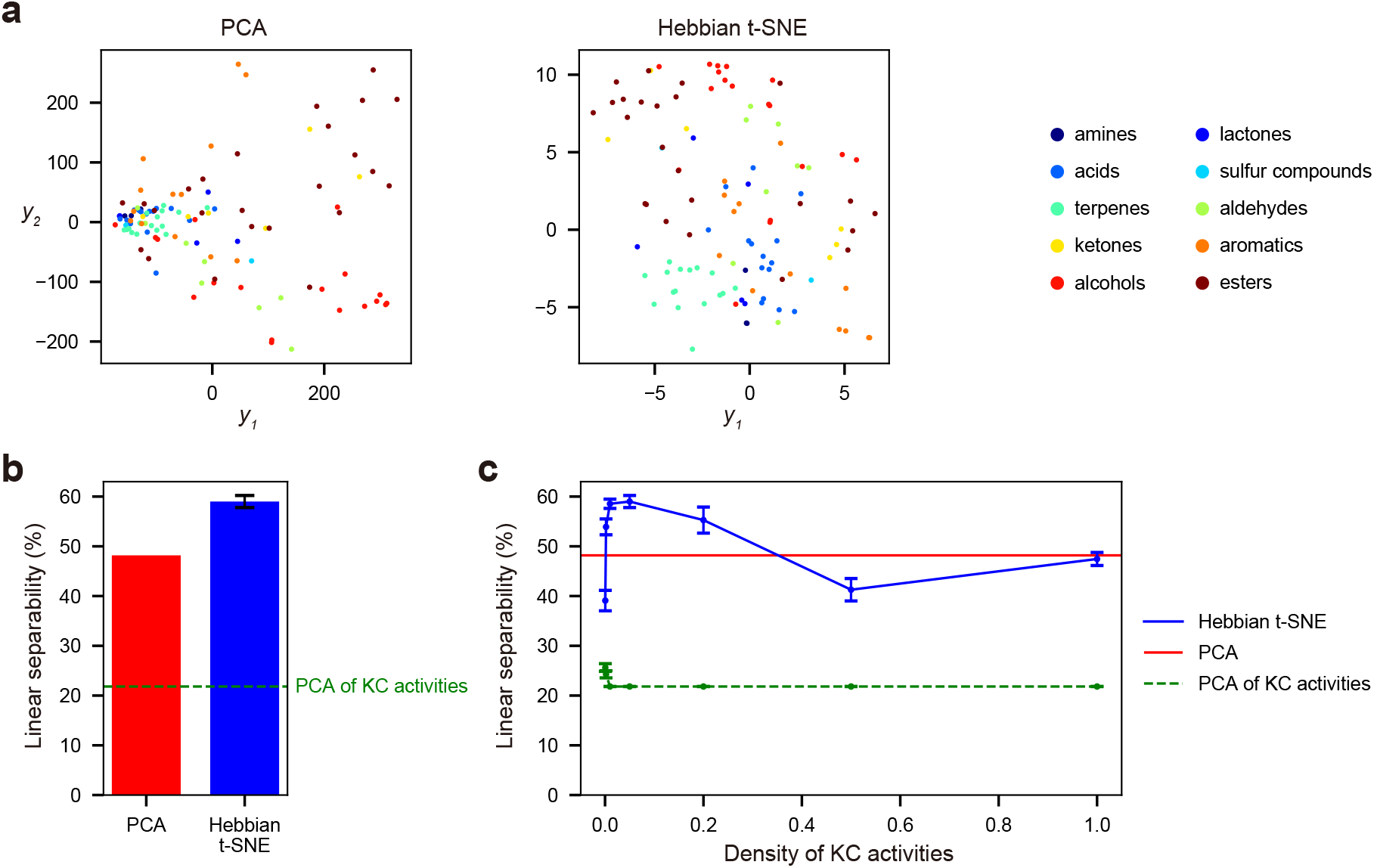
The Hebbian t-SNE obtained a good chemical representation behind olfactory response in Drosophila. (a) The representations obtained by applying the PCA (left) and Hebbian t-SNE (right) to olfactory receptor responses. The colors indicate the ten functional groups (amines, lactones, acids, sulfur compounds, terpenes, aldehydes, ketones, aromatics, alcohols, and esters) based on the chemical structures. (b) The linear separability of the ten functional groups in the two-dimensional representations obtained by the PCA and Hebbian t-SNE. The error bar indicates SEMs in the ten trials. The green dotted line indicates the linear separability obtained by applying the PCA to the modeled KC activities. (c) The linear separability of the ten functional groups based on the two-dimensional output representations as the density of KC activities was varied. The error bars indicate SEMs in the ten trials.

### Hebbian t-SNE might be working in the olfactory circuits in Drosophila

Here, we examined the potential role of the Hebbian t-SNE in biological circuits. The Hebbian t-SNE requires the middle layer that is composed of a large number of neurons with sparse neuronal activity. This structure is observed in the olfactory circuit in Drosophila, comprising PNs, KCs, and MBONs [18]. Thus, we modeled the PNs, KCs, and MBONs as the input layer *X*, middle layer *Z*, and output layer *Y*, respectively (Fig. 1b), and investigated whether the Hebbian t-SNE might work during olfactory learning in Drosophila.

We first analyzed the responses of 24 olfactory receptors to 110 odors, which are classified into ten functional groups based on the chemical structures [19]. Considering that chemically-related odors are reported to elicit similar neuronal representations and perceptions [27, 28, 29], we evaluated whether the PCA and Hebbian t-SNE can find the representation reflecting this chemical classification. As olfactory receptor neurons expressing the same olfactory receptor transmit signals to identical PNs [30, 31, 32], we utilized these receptors’ responses as surrogates of PN activities. First, the representations applying the PCA to the olfactory receptor responses crudely distinguished different chemical functional groups (Fig. 4a). In Drosophila, the transformation from the PN to KC activities is reported to be described as a combination of the random projection and lateral inhibition [33, 34]. Therefore, we estimated the KC activities by applying random projection, thresholding, and normalization for the PN activities (see Methods), consistent with previous studies [33, 35, 36, 37]. We conducted Hebbian t-SNE employing these estimated KC activities as the activities of the middle layer. As a result, the Hebbian t-SNE obtained representations distinguishing the different chemical functional groups better than the PCA of olfactory receptor responses (Fig. 4a). The linear separability of different chemical classes was significantly higher in the Hebbian t-SNE (59 %) than in the PCA (48 %) (Fig. 4b). This good representation by the Hebbian t-SNE was observed when the density of the KC activities was low (Fig. 4c). However, excessively sparse KC activities hindered proper discrimination since the KC activities for some inputs completely overlapped in this case. With KC representations that are too densely overlapping, independent learning of the output response to each input became impossible. Therefore, the intermediate density, including the experimentally observed level of 5 % [18], was optimal for linear separability across different patterns. Note that the representations by applying the PCA to the modeled KC activities were unsuccessful since the input structure was distorted by the random transformation from PN to KC activities (Fig. 4b and c). These results were robust to changes in the perplexity value (Extended Data Fig. 4). Hence, the Hebbian t-SNE effectively captures the chemical structure behind high-dimensional olfactory sensory inputs, which implies its benefit for downstream olfactory processing.

Next, we investigated whether the representation by the Hebbian t-SNE can extract valence, which is expressed in the MBON activities of Drosophila [17]. For this purpose, we analyzed the experimental data from [20], where the responses of 37 types of PNs to 84 odors were recorded together with the valence index, which evaluates how much flies prefer the odor by monitoring their behavioral responses. While this study did not record the activities of MBONs, the valence index is expected to be encoded in MBON activities because the previous study [17] found that odorants sharing similar valence index were clustered within the low dimensional representation of MBON activities. Hence, we analyzed whether the t-SNE could obtain a representation consistent with this valence index. We applied the Hebbian t-SNE to the responses of 37 types of PNs and compared the result with the PCA representations of the PN activities and the estimated KC activities, computed as in Fig. 4 (Fig. 5a). While the representations were only weakly correlated with the valence index when evaluated in the whole range (Extended Data Fig. 5), nearby points tended to have a more similar valence index in the representation obtained by the Hebbian t-SNE than the PCA representations of the PN and modeled KC activities. For an objective assessment, we performed the K-means clustering on each two-dimensional representation. We compared the three representations by the sum of the squared error between experimentally measured valence index values and their group mean within each cluster (Fig. 5b). The representation by the Hebbian t-SNE was locally more homogeneous in the valence index than the PCA representations of the PN and modeled KC activities (Fig. 5c). Given that odors with similar valence tend to cluster in the MBON representation [17], this result suggests that the Hebbian t-SNE might be the most plausible computation conducted in this circuit among the three methods. The outcomes were robust to different choices of the perplexity value (Extended Data Fig. 6). These results indicate the possibility that nonlinear dimensionality reduction, such as the Hebbian t-SNE, can be implemented in Drosophila olfactory circuits.

**Figure 5.**
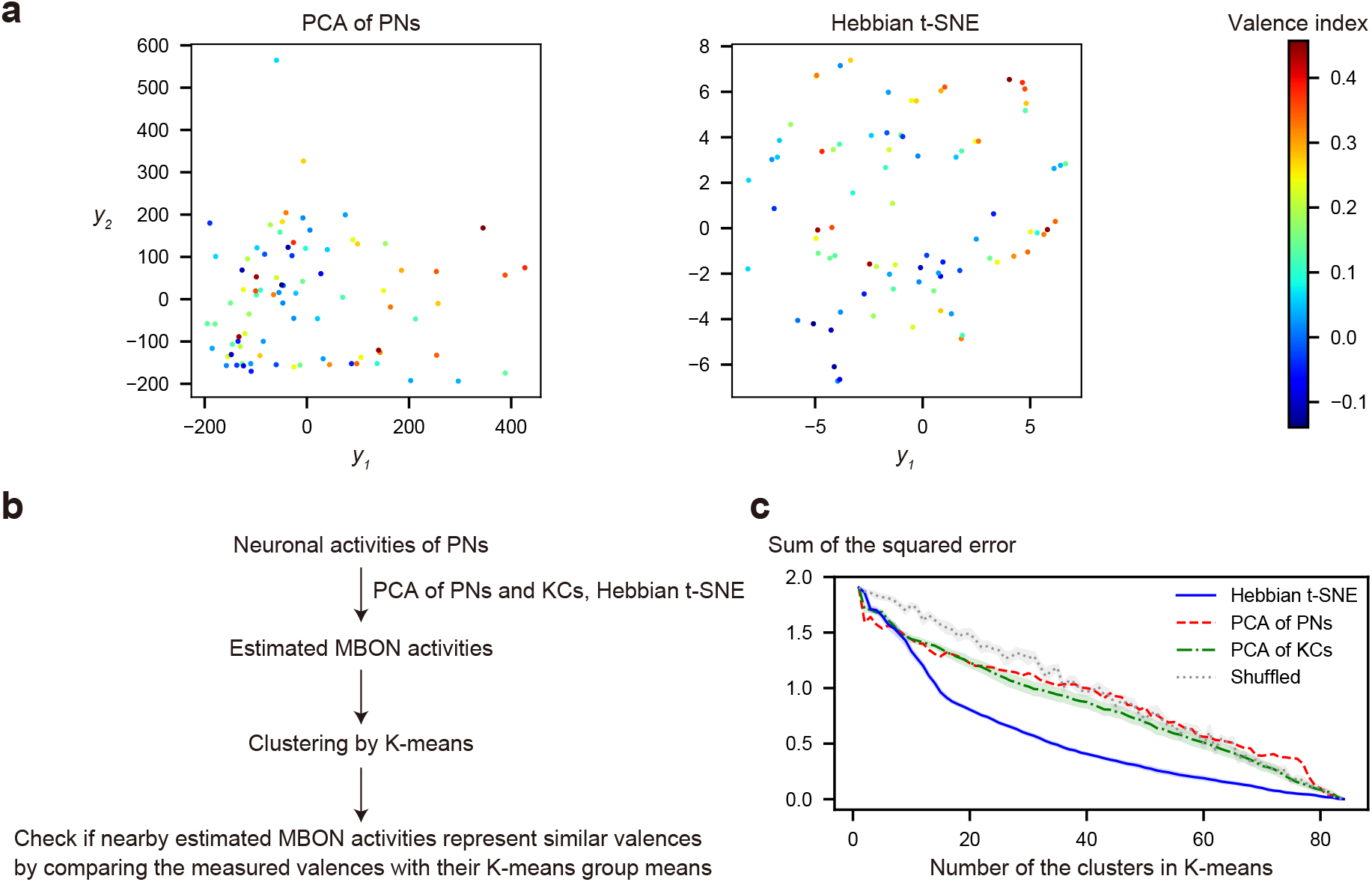
The Hebbian t-SNE explained valence representation in Drosophila better than the PCA. (a) The two-dimensional representations by applying the PCA (left) and Hebbian t-SNE (right) to the PN activities. The colors indicate the valence index. (b) The procedure of the analysis. (c) The squared errors between the valence index of each data point and its neighborhood cluster mean for varying numbers of K-means clusters. The representations were obtained by applying the PCA to the PN activities (red), PCA to the modeled KC activities (green), and Hebbian t-SNE to the PN activities (blue). The gray line indicates the result of Hebbian t-SNE but after random shuffling of the valence index. The lines and shadows represent the means and SEMs in the ten trials, respectively.

### Hebbian t-SNE induces association learning with generalization

The MBON representations are not formed entirely in an unsupervised manner. In Drosophila, the association between odors and rewards is detected by DANs, whose activities promote synaptic plasticity from KCs to MBONs [17, 38, 39]. Additionally, a recent study reported that DANs are also activated by the innate value of each odor [40]. Thus, we explored the collaborative role of the Hebbian t-SNE in conjunction with partially available rewards and the innate value of odors. For simplicity, we subsequently use the term ‘rewards’ to denote the effects of both the externally provided rewards and the innate value of odors. Mimicking the synaptic plasticity mediated by DANs, we considered a simple rule in which the reward associated with odors induces modification of synaptic weight from the middle layer to a subset of the output neurons (*y*_1_ in the case below) in addition to synaptic changes by the Hebbian t-SNE. This corresponds to adding a reward-modulation term in the objective function of the t-SNE (see Methods). Thus, the total synaptic changes are computed using 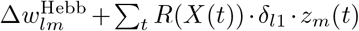 where 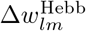 is a synaptic change by the Hebbian t-SNE, *R*(*X*(*t*)) represents the reward associated with the input *X*(*t*), *δ*_*l*1_ is Kronecker delta function that restricts changes only in synapses to *y*_1_, and the *z*_*m*_(*t*) factor induces synaptic changes only when presynaptic neuron *m* is active (see Methods for details). Note that, biologically, positive associations are often implemented by the depression of synapses onto the MBONs associated with avoidance behavior in Drosophila [18, 38, 41]. These neurons’ activity might be aligned with the negative-*y*_1_ direction in our model. Given that the MBON activities predict attraction and repulsion behaviors to odors [42], we investigated whether this reward-modulated Hebbian t-SNE could obtain the representation separating positively and negatively rewarded inputs.

We considered input data composed of four entangled rings (Fig. 6a). We applied the reward-modulated Hebbian t-SNE in the case that positive and negative rewards were given to a subset of inputs within each ring (Fig. 6a). Two rings were associated with positive rewards, and the others were associated with negative rewards. As a result, the reward-modulated Hebbian t-SNE separated the positive-rewarded and negative-rewarded rings even when only a minority of inputs were paired with reward signals (Fig. 6b). When the reward proportion was set to 0.1, the reward-modulated Hebbian t-SNE obtained a representation similar to the case with reward proportion 1.0 (Fig. 6b). This is qualitatively evaluated using a reward separability, which assesses the separability of the rings associated with positive and negative rewards based on their values of *y*_1_ (Fig. 6c; see Methods for a definition). The reward separability improved significantly even with a small reward proportion *∼* 0.1 (Fig. 6d). Thus, the reward signal was generalized to the inputs that were not rewarded but belonged to the same ring as rewarded inputs. This is consistent with the olfactory learning in Drosophila in which avoidance is generalized among similar odors [43].

**Figure 6.**
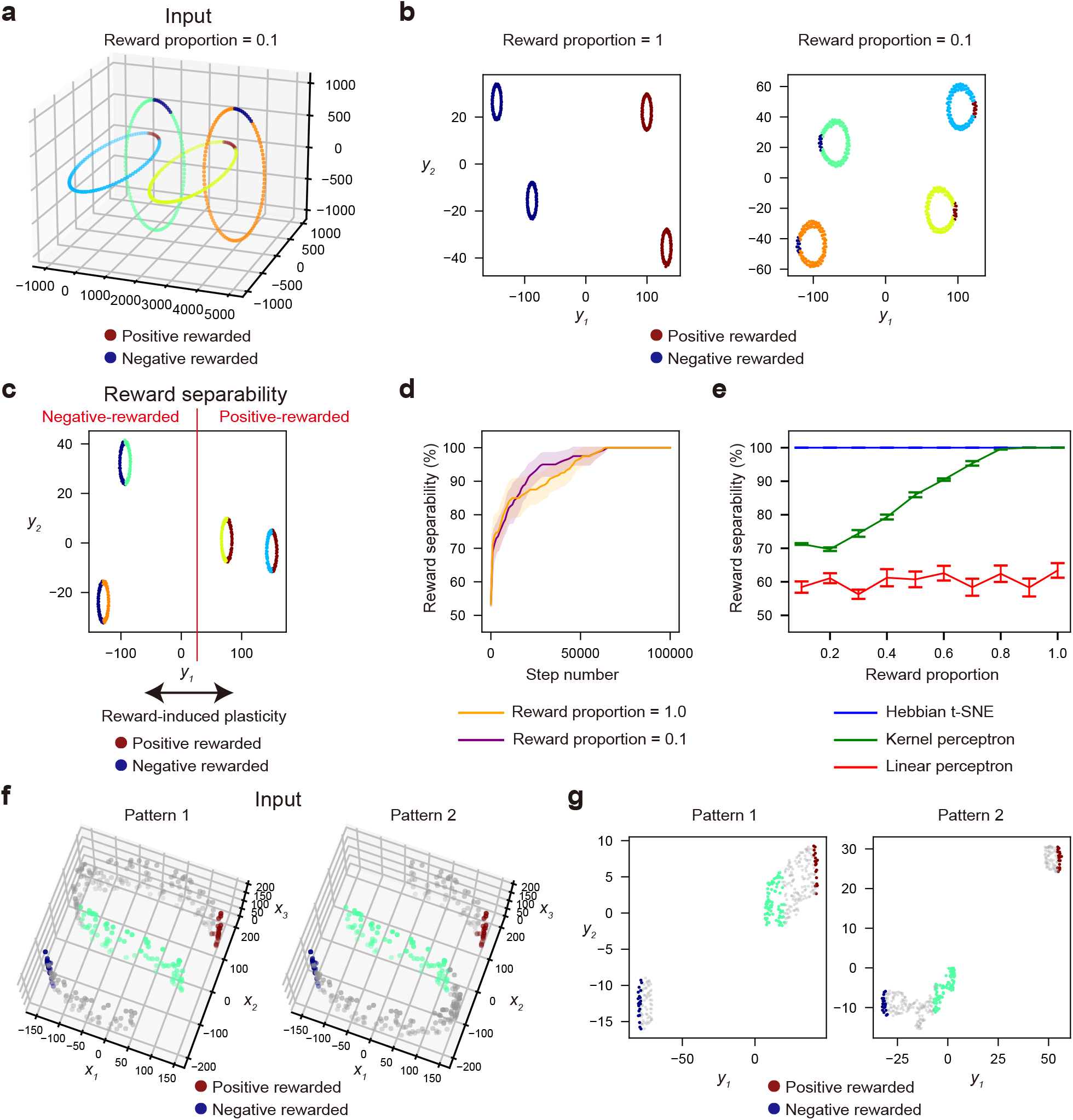
Reward-modulated Hebbian t-SNE. (a) The original data of the four entangled rings in the three-dimensional space. The red and blue points indicate positive- and negative-rewarded data, respectively. The reward proportion is the proportion of the rewarded data in the total data. (b) The representations obtained with distinct reward proportions. The red and blue data points accompanied rewards. Their representations were biased to the positive and negative direction of the *y*_1_ coordinate by the positive and negative reward, respectively. (c) Reward separability is defined as the classification accuracy of whether each data point belongs to a positively or negatively rewarded ring based on output *y*_1_ when its threshold is optimally chosen. The reward proportion was 0.5 in this example. (d) The time course of reward separability when the reward proportion was 1.0 (orange) and 0.1 (purple). The lines and shadows represent the means and SEMs in the ten trials, respectively. (e) The reward separability of the Hebbian t-SNE (blue), linear perceptron for three-dimensional inputs (red), and linear perceptron after transforming inputs into high-dimensional space (green) (see Methods) for different reward proportions. The error bars represent SEMs in the ten trials. (f) Three-dimensional input patterns along disconnected S-shaped sheets, a part of which is cut. The red and blue points indicate positive- and negative-rewarded data, respectively. The number of the rewarded points is equal to that in Extended Data Fig. 7a at reward proportion 0.1. The light green points are some neutral data points located around the center of the S-shape, which are colored for visualization purposes. The manifolds are cut so that the light green data are connected only to the positive-rewarded data in pattern 1 and the negative-rewarded data in pattern 2. (g) The representations obtained by the reward-modulated Hebbian t-SNE. The light green points were located near the positive- and negative-rewarded data in patterns 1 and 2, respectively.

To investigate the generalization ability further, we compared the reward-modulated Hebbian t-SNE with linear and kernel perceptrons. Systematically changing the reward proportion suggested that the Hebbian t-SNE had a higher generalization ability than these perceptrons (Fig. 6e). The simple linear perceptron, given the three-dimensional inputs (see Methods), could not separate the rings associated with positive and negative rewards because they were entangled. The kernel perceptron used linear perceptron after transforming the original inputs to a high-dimensional representation using the Gaussian kernel (see Methods). This kernel perceptron had better performance than the linear perceptron. Yet, the Hebbian t-SNE had superior generalization ability to this method (Fig. 6e). While the kernel perceptron only incorporates the distance from each point to the rewarded points, similar to the nearest neighbor algorithm, the Hebbian t-SNE is expected to have a stronger generalization ability utilizing geometric structures in the original inputs. We also considered the data along an S-shaped manifold, in which one end is positively rewarded, and the other is negatively rewarded (Extended Data Fig. 7). In this case, the reward-modulated Hebbian t-SNE also had superior generalization ability to the other two methods. Furthermore, when the S-shaped manifold had discontinuity towards the negative- or positive-rewarded data (Fig. 6f), the data around the center of the S-shaped curve acquired either positive or negative valence, respectively (Fig. 6g), reflecting the geometric continuity along the S-shaped manifold. This result suggests that the generalization by the reward-modulated Hebbian t-SNE depends on the distribution of whole input patterns, including the non-rewarded data. This can be experimentally tested by providing different distributions of odor mixtures of positive-rewarded and negative-rewarded odors in Drosophila (see Discussion). In summary, the reward-modulated Hebbian t-SNE facilitates association learning with generalization ability.

## Discussion

We showed that the Hebbian t-SNE, constructed by the simple feedforward neuronal network with the Hebbian synaptic plasticity learning rule, performed comparably to t-SNE on datasets including the entangled rings and MNIST data. We then suggested the possibility that the Hebbian t-SNE might be implemented for learning low-dimensional representations of the high-dimensional odor space in Drosophila. We further showed that the reward-modulated Hebbian t-SNE enabled association learning with generalization, as with the olfactory systems in Drosophila. These findings suggest that even simple biological circuits can conduct nonlinear dimensionality reduction and obtain untangled representations in an unsupervised way.

The proposed model is based on previous experimental findings in neuroscience, especially of the network architecture and synaptic plasticity rule in the olfactory circuit in Drosophila. The model proposes multiple essential characteristics, the role of which could be tested in future experiments. First, high-dimensional activities must be created by a large number of neurons in the middle layer. In Drosophila, KCs are experimentally reported to exhibit higher-dimensional representations of odors compared to PNs [33]. Similar to the Drosophila olfactory circuit, a comparable neuronal circuit has been identified in the cerebellum, including a large number of granule cells in the middle layer [37, 18]. In our model, inputs are transformed into high-dimensional space in the middle layer, allowing output representations to move independently. Therefore, the number of middle-layer neurons influences the precision for distinguishing similar data. Since very similar inputs (e.g., similar odors) need not necessarily be distinguished in many cases, the number of neurons in the aforementioned biological circuits (*∼* 2000 KCs in Drosophila and *∼* 50*−*70 billion cerebellar granule cells in humans [44]) might be sufficient to achieve representations with enough precision. Our model predicts that intermediate sparseness is optimal for achieving high-dimensional activities in a fixed number of middle-layer neurons (Fig. 4c). The sparseness of KC activities is known to be modulated by inhibitory neurons called anterior paired lateral (APL) neurons [34]. Thus, experimentally modulating APL neurons could test the model prediction.

Second, the computation of the global factor *D* plays a critical role in this model. The proposed synaptic plasticity rule is written in the form of the three-factor learning rule [45, 46], composed of the presynaptic activity, postsynaptic activity, and the global factor. Specifically, the presynaptic and postsynaptic terms are described as the difference in their activities at neighboring time points. This formulation aligns with previously proposed differential Hebbian learning, potentially corresponding to experimentally observed spike-timing-dependent plasticity in spiking neurons [47, 48]. In addition to the presynaptic and postsynaptic activities, the global factor *D* regulates network outcomes by incorporating common information among synapses. In the example of the olfactory circuit in Drosophila, one suggested global factor is the activity of DANs, which is known to modulate the synaptic plasticity from KCs to MBONs [38, 40, 49]. Anatomically, the neurons corresponding to 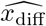 and 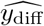 which project to DANs might be specific types of KCs and MBONs [50, 51]. Although the activities corresponding to 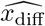 and 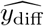 have not yet been reported to our knowledge, similar components that calculate the difference between the current neuronal activity (*X*(*t*) and *Y* (*t*)) and previous neuronal activity (*X*(*t −* 1) and *Y* (*t−* 1))), might be computed using a neuronal adaptation mechanism. One study suggested that the flies’ responses to odors showed a relatively slow adaptation over a timescale of several minutes [20]. Therefore, it might be possible that the adaptation at a timescale compatible with the Hebbian t-SNE also exists and calculates the time derivative of neuronal activities. The suggested role of the DANs in this model (i.e., comparing the input and output similarities) is a new prediction to be further investigated in the future.

The third point is the biological mechanism by which inputs are repeatedly presented in a random order in our model. One possibility is that such a process can be realized by experiencing real inputs many times. A recent study suggested that odor exposure without rewards can cause synaptic plasticity [40]. In the future, changes in representation resulting from such passive odor exposures could be experimentally compared with the predictions of our model. Another possibility is to exploit offline learning with memory reactivation [52, 53, 54, 55]. During rest and non-rapid eye movement (NREM) sleep, neural activities during awake experiences are reported to be reactivated [24, 56, 57]. Repeated neuronal reactivation might be utilized to reorganize memory, as our model suggests. Representation learning during rest and sleep periods is an interesting future topic.

In this study, the reward-modulated Hebbian t-SNE conducts association learning with generalization ability. The generalization can also be realized by the overlap of the KC activities. A previous study reported that flies’ responses are generalized to similar odors, which share the KCs’ activities with the original odor [38]. In this case, the generalization happens simply because the two odors share the synapses from KCs to MBONs. This overlap of KCs between similar odors is an important aspect related to locality-sensitive hashing used in similarity searches [35, 58]. On the one hand, the Hebbian t-SNE also works in the case that the representations in the middle layer *Z* overlap with each other (Figs. 4-5 and Extended Data Figs. 4-6). On the other hand, our study suggests the complementary possibility that generalization can occur even without shared KC activities, utilizing the input geometric structures. In Fig. 6, we suggested that the reward-modulated Hebbian t-SNE has superior generalization ability to the kernel perceptron. The generalization capability of the kernel perceptron exclusively relied on the overlapped KC activity in the model. Therefore, these results suggested an additional benefit of the Hebbian t-SNE utilizing input geometric structures complementary to the KC activity overlaps. Crucially, the contribution to the generalization by the Hebbian t-SNE independent of the KC activity overlap can be tested by the experiment using odor mixtures in Drosophila (Fig. 6f and g). Our model predicts that, when one odor A is associated with reward and another odor B is associated with punishment, the valence of the 50% mixture of the two odors should depend on the continuity of odor mixtures used for learning. The acquired valence of the 50% mixture in this model differs depending on whether a mixture gap segregates positively or negatively rewarded odors (Fig. 6g), even when the KC activity overlap is identical. Thus, these two different generalization strategies can be experimentally tested in the future.

One study reported that conducting the dimensional reduction with the t-SNE before providing the data in the convolutional neural network (CNN) is useful for improving performance [59]. Although this study assessed the representation by the linear separability in line with the previous studies, e.g., [60]), the obtained representation by the Hebbian t-SNE could also be useful for more complex downstream models such as CNNs. Additionally, the Hebbian t-SNE could be further extended for biological realizations of representation learning methods related to contrastive learning [61]. Contrastive learning provides representations that are not necessarily low-dimensional in a self-supervised manner. Some recent studies proposed biologically plausible algorithms related to contrastive learning in deep neural networks [9, 62]. They are based on the idea that similar inputs should also be close in the output space, and can be described within the same framework as t-SNE [63, 64]. Therefore, our setup might be extended to useful representation learning beyond dimensionality reduction in a simple architecture with a biological-plausible learning rule.

In summary, our model suggests that simple biological circuits with Hebbian synaptic plasticity can conduct nonlinear dimensionality reduction.

## Methods

### Hebbian t-SNE

In this section, we concisely introduce the Hebbian t-SNE algorithm. See the next “Comparison between the t-SNE and Hebbian t-SNE” section for how this algorithm is derived based on the original t-SNE.

Let us consider a three-layer neural network model. The three layers were denoted by *X* = (*x*_1_, *x*_2_, …, *x*_*r*_), *Z* = (*z*_1_, *z*_2_, …, *z*_*s*_), and *Y* = (*y*_1_, *y*_2_). The whole *N* patterns *X*^1^, *X*^2^, …, *X*^*N*^ were presented in a random order numerous times. The values of *X, Z*, and *Y* at time *t* were denoted by *X*(*t*) = (*x*_1_(*t*), *x*_2_(*t*), …, *x*_*r*_(*t*)), *Z*(*t*) = (*z*_1_(*t*), *z*_2_(*t*), …, *z*_*s*_(*t*)), and *Y* (*t*) = (*y*_1_(*t*), *y*_2_(*t*)) (*t* = 0, 1, 2, …), respectively. The patterns of neighboring epochs were assumed to be different, thus *X*(*t*)*≠X*(*t−* 1). The middle layer activities *Z* in response to different input patterns were required to be linearly independent for an accurate implementation of the t-SNE. Except for Figs. 4-5 and Extended Data Figs. 4-6, the transformation from *X* to *Z* was set as a winner-take-all. The projection from *Z* to *Y* was linear, thus *Y* (*t*) = *W* (*t*)*Z*(*t*), where *W* (*t*) = (*w*_*lm*_(*t*)) is a synaptic weight matrix. We considered a learning rule of synaptic weight *W*, inspired by the t-SNE. We adopted a batch update with *T* steps to speed up the simulation. The synaptic weights were updated with Adam [25], described as

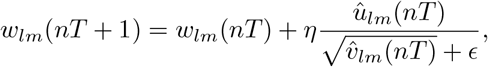

with η = 0.1 and ϵ = 10−8. The terms 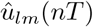and 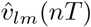were described as

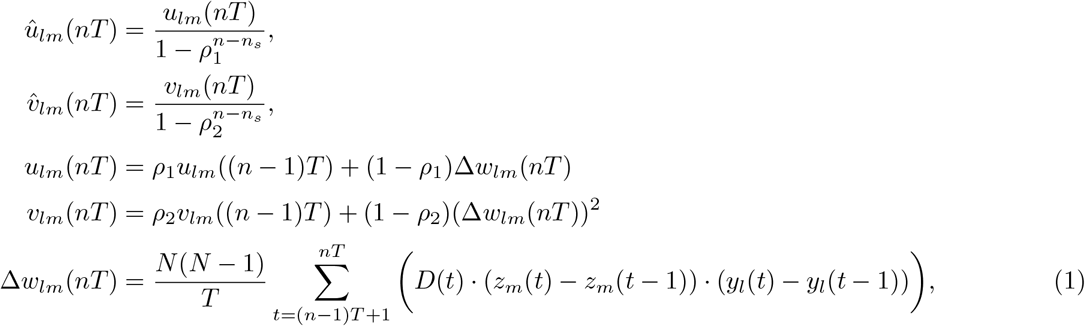

with *ρ*_1_ = 0.9 and *ρ*_2_ = 0.999. Note that the factor Δ*w*_*lm*_(*nT*) represents the gradient of the loss function in Eq. (6). Synaptic weights were changed after the initial warmup steps (*n > n*_*s*_ with *n*_*s*_ = 500), when the perplexity approached sufficiently close to the target value (Figs. 2e and 3e). The initial values *u*_*lm*_(*n*_*s*_*T*) and *v*_*lm*_(*n*_*s*_*T*) were set to zero.

Next, we describe how the global factor *D*(*t*) was computed. The neuronal activity corresponding to *D*(*t*) received inputs 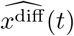 and 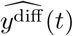 (see Fig. 1 and Table 1), which estimate input and output similarities, respectively. The neuronal activity 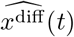 was calculated using the middle layer activity *Z* and the input change ∥*X*(*t*) *− X*(*t −* 1)∥^2^. First, the axonal activity *a*_*k*_(*t*) based on the middle layer activity *z*_*k*_(*t*) was set to one at time *t* when the *z*_*k*_(*t*) exhibited the largest positive value among the responses to the input patterns *X*^1^, *X*^2^, …, *X*^*N*^, and otherwise *a*_*k*_(*t*) = 0. This mechanism can be biologically implemented, for example, by adjusting each neuron’s firing threshold to achieve an average firing rate of 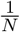 (In non-generic cases, where *z*_*k*_(*t*) took the largest value for multiple input patterns, *a*_*k*_(*t*) was set to one for a randomly selected input pattern among them and zero otherwise.) Subsequently, each synapse had a synapse specific variable *σ*_*k*_ and a postsynaptic weight 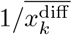. The transmitter release of the synapse *k* was determined by 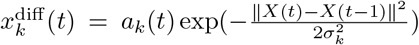 Finally, the postsynaptic neuronal activity 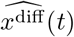 was described using the transmitter release 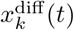 and the postsynaptic weight 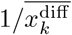 as

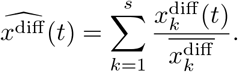

The postsynaptic weight 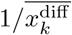 were batch-updated so that the 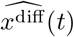 was appropriately normalized according to

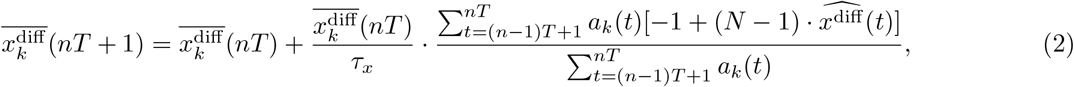

for *n ≥* 1 with *τ*_*x*_ = 100. This normalization makes sure the contribution from each pattern is equal regardless of the number *K* of simultaneously active axons (See the next “Comparison between the t-SNE and Hebbian t-SNE” section for details). The variable *σ*_*k*_ was regulated to achieve the desired perplexity of the t-SNE by introducing *H*_*k*_(*t*), which estimates information entropy computed as

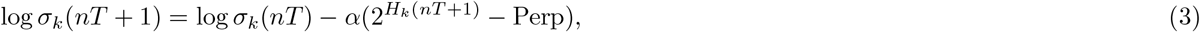

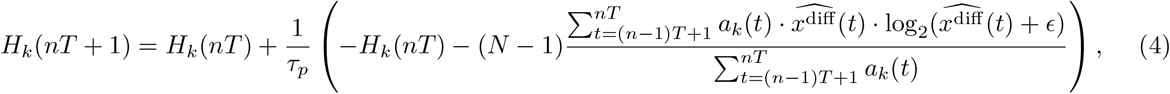

for *n ≥* 1, with *τ*_*p*_ = 100, *α* = 10^*−*3^, and *ϵ* = 10^*−*8^, which prevents the logarithm from being imaginary. The term Perp corresponds to the target perplexity in the t-SNE, which was set to 20 in Figs. 2, 4-6, and Extended Data Fig. 5, 40 in Fig. 3 and Extended Data Fig. 7, and 30 in Extended Data Figs. 4 and 6.

The neuronal activity 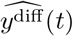 was calculated using *y*^diff^ (*t*) as

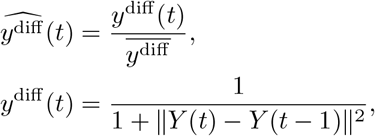

where the time average 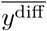 was batch-updated as

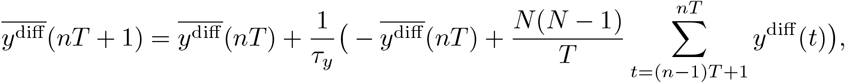

for *n ≥* 1, with time constant *τ*_*y*_ = 100. Finally, the neuronal activity *D*(*t*) was calculated as

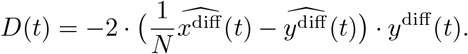

The variables *w*_*lm*_(*t*), 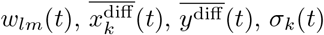, *σ*_*k*_(*t*), and *H*_*k*_(*t*) were equal to the values of time *t −* 1 except for the case described above.

### Comparison between the t-SNE and Hebbian t-SNE

In this section, we derive the Hebbian t-SNE algorithm based on the t-SNE [15]. First, we briefly review the original t-SNE. In the t-SNE, the similarity *p*_*ij*_ between inputs *X*^*i*^ and *X*^*j*^ is defined as

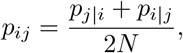

where *N* denotes the total number of inputs and *p*_*i*|*j*_ is described as

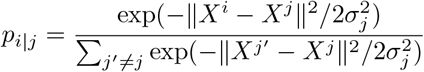

for *i≠j* and *p*_*i*|*i*_ = 0 for all *i*. The variables *σ*_*j*_ are determined by searching the value that achieves a suitable perplexity specified by the user for each data point *j*. The perplexity of data point *j* is defined as

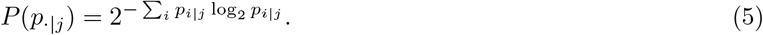

The similarity *q*_*ij*_ of outputs *Y* ^*i*^ and *Y* ^*j*^ are described as

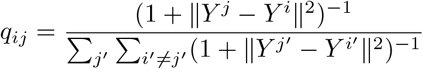

for *i≠j* and *q*_*i*|*i*_ = 0 for all *i*. The t-SNE aims to minimize the Kullback-Leibler divergence between the input similarities *p*_*ij*_ and the output similarities *q*_*ij*_. Thus, the objective function *C* is described as

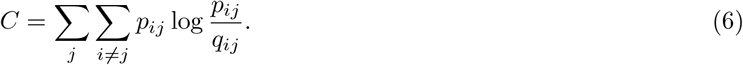

The gradient of this function *C* is given by

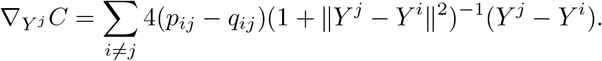

Therefore, the mapping to output *Y* is changed according to this gradient. Note that *Z*_*j*_ in Table 1 is a column vector whose *j*-th component is one and the other components are zero. If the mapping to output is represented as *Y* ^*j*^ = *WZ*^*j*^, then the gradient with respect to the synaptic weight matrix *W* = (*w*_*lm*_) can be described as

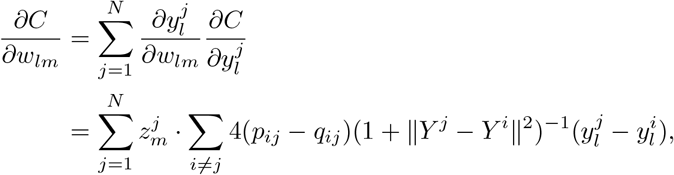

where 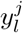 and 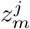 are the *l*-th and *m*-th components of *Y* ^*j*^ and *Z*^*j*^, respectively.

In the following paragraphs, we prove that the expected value of the synaptic changes in the Hebbian t-SNE Δ*w*_*lm*_(*nT*) in Eq. (1) is proportional to the negative gradient 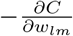 under the condition that the batch size *T* is sufficiently large and the time-average variables 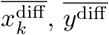, and *H*_*k*_ are calculated over sufficiently long time. From here, *A → B* indicates that the value *A* approaches to the value *B* under this condition.

We first consider the case of *X*(*t*) = *X*^*j*^ and *X*(*t −* 1) = *X*^*i*^. Let *k*_1_, *k*_2_, …, *k*_*K*_ be a set of all indices *k* such that *a*_*k*_(*t*) = 1 in this case. Since each axon *k* responds only to a single input pattern, the axonal activities 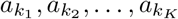 take positive value only when the input pattern is *X*^*j*^. Moreover, since 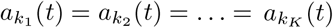 at all time *t*, the values of 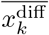 and *σ*_*k*_ are equal for all *k* = *k*_1_, *k*_2_, …, *k*_*K*_. Therefore, we obtain

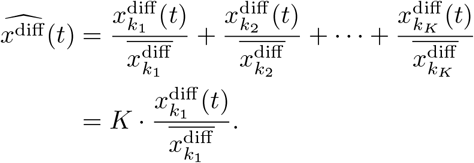

By using this formula, the update of 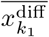 in Eq. (2) is described as

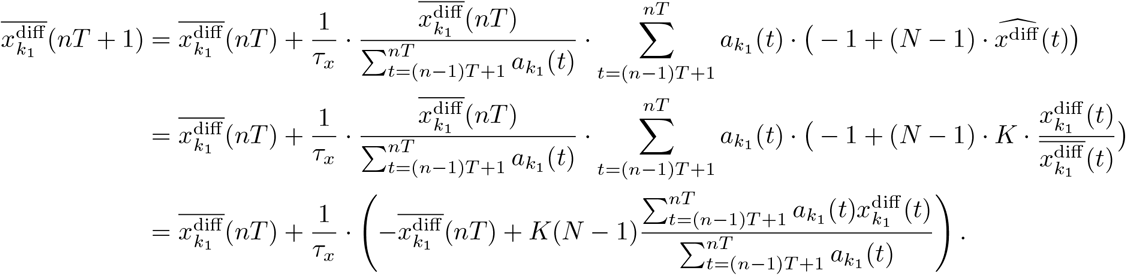

The last equality is followed by 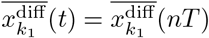 for *t* in (*n −* 1)*T* + 1 *≤ t ≤ nT* because of the batch update. Therefore, noting 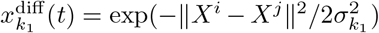 when *X*(*t*) = *X*^*j*^ and *X*(*t−* 1) = *X*^*i*^, we obtain after many iterations

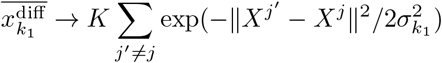

because of the low of large numbers and because every pattern pair appears with equal probability over the duration *T*. By using these formulas,

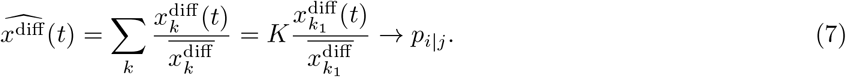

Note that *σ*_*k*_1 achieves a desired perplexity in the same way as with the t-SNE for the following reason. From Eq. (7), we find

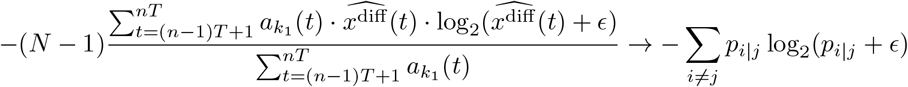

Following Eq. (4), the *H*_*k*_1 (*t*) approaches the left-hand side of this equation. Hence,

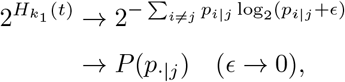

where *P* (*p*_*·*|*j*_) represents the perplexity of data point *j* in Eq. (5). Since the 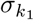 is regulated so that 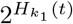 approaches the target perplexity in Eq. (3), 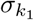 achieves the desired perplexity in the same way as the t-SNE.

Also,

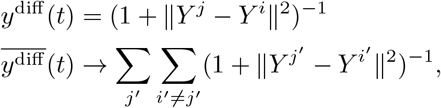

which leads to 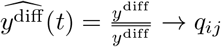 Therefore, using the definition of *D*(*t*) and *D*_*i*|*j*_ in Table 1,

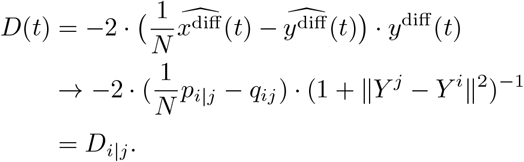

Then, averaging over *t* with random pattern presentations, we obtain

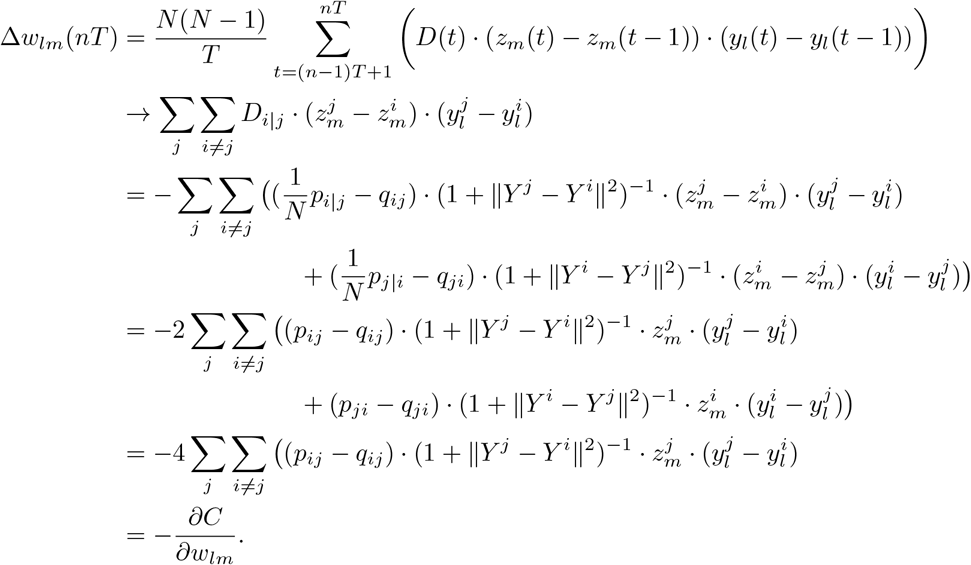

In summary, the proposed algorithm realizes t-SNE in the biological neural circuit.

### Simplified Hebbian t-SNE

The only difference from the original model was in the formulation of 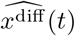. In the simplified model, the term 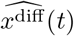 was described as

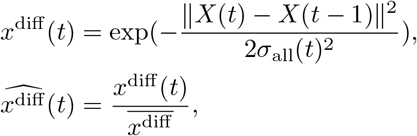

where the variable 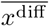 was updated by

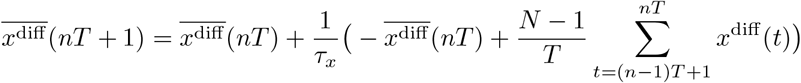

for *n ≥* 1, with time constant *τ*_*x*_ = 100. The variance *σ*_all_ common to all input patterns, was updated by

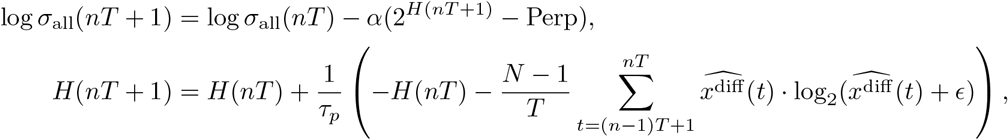

for *n ≥* 1, with *τ*_*p*_ = 100, *α* = 10^*−*3^, *ϵ* = 10^*−*8^, which prevents the logarithm from being imaginary, and Perp corresponds to the target perplexity in the t-SNE, which was set to 20 in Extended Data Fig. 2 and 40 in Extended Data Fig. 3.

### Reward-modulated Hebbian t-SNE

We introduced a reward-modulation term into the objective function as follows: *L* = *C − λC*_R_ with *λ* = 1 and 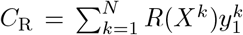, where *R*(*X*^*k*^) represents the reward associated with the input *X*^*k*^. The gradient was described as

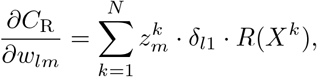

where *δ* is Kronecker delta. By defining 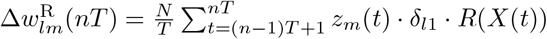 then

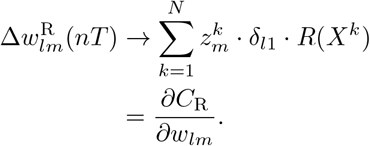

Therefore, we added the 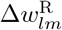 to the Hebbian t-SNE as a reward modulation as follows:

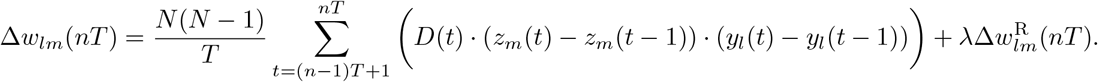

In Fig. 6 and Extended Data Fig. 7, the reward *R*(*X*(*t*)) was set to 7.5 *·* 10^*−*6^*/p* and *−*7.5 *·* 10^*−*6^*/p* for positive- and negative-rewarded input patterns, respectively, where *p* denotes the proportion of rewarded patterns. The reward *R*(*X*(*t*)) was set to zero for other non-rewarded input patterns.

### Linear methods with transformed high-dimensional data

In Fig. 6 and Extended Data Fig. 7, the original three-dimensional input patterns were transformed into high-dimensional activities *Z* = (*z*_1_, *z*_2_, …, *z*_*N*_), where *N* is the total number of input patterns. Each original input *X*^*j*^ was transformed into 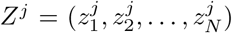 with 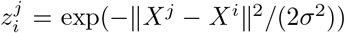. The optimal *σ* value was searched among 5, 10, …, 1000 and determined to maximize the average reward separability across the reward proportions ranging from 0.1 to 1.0. Consequently, *σ* was set to 25 in Fig. 6 and 915 in Extended Data Fig. 7. In Fig. 6 and Extended Data Fig. 7, the linear perceptron, the linear perceptron adapted for transformed high-dimensional data, and the Hebbian t-SNE were compared. The generalization performance of the linear perceptron was calculated as follows. The linear perceptron, implemented with the scikit-learn package [65] in Python, was trained only using the positive- and negative rewarded points. All input patterns were then projected into a one-dimensional space using the obtained weight of the linear perceptron. The reward separability of the projected one-dimensional value was calculated.

### Estimation of the KC activities

In Figs. 4-5 and Extended Data Figs. 4-6, the KC activities *Z* = (*z*_1_, *z*_2_, …, *z*_2000_) were estimated by using a theoretical model composed of random projection, thresholding, and normalization, consistent with the previous studies [33, 35]. We constructed the random weight matrix *W*_rand_ from the PNs *X* to KCs *Z*, in which each KC received inputs from seven randomly chosen PNs and those synaptic weights were set to one. Since the other synaptic weights were set to zero, the sum of each row of *W*_rand_ was seven. For a given PN activity *X*^*j*^, the corresponding *Z*^*j*^ was defined as follows. First, 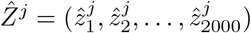 was computed as

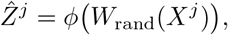

where the function *ϕ* is a threshold-linear function that outputs the original value when the input is positive and is among the largest *p* proportion of the total 2000 neurons and zero otherwise. The density *p* of the KC activities was set to 0.05 according to previous experimental observation [18] (but explored for various densities in Fig. 4c and Extended Data Fig. 4c). Then, the estimated KC activity *Z*^*j*^ was obtained by normalizing 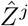 described as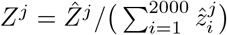. The thresholding and normalization can be realized through global inhibition by APL neurons in the Drosophila olfactory circuit [34].

### Other Simulation details

The initial value of each component in the weight matrix *W* was randomly sampled from the standard Gaussian distribution. The initial value of *σ*_*k*_ was set to 500. The initial values of 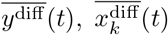, and *H*_*k*_(*t*) were described as

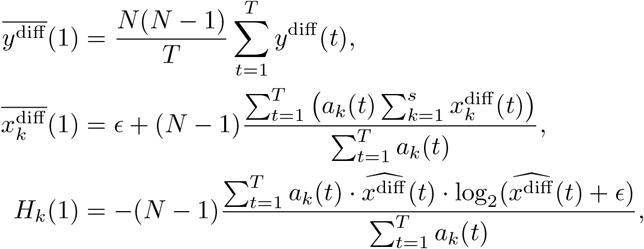

with *ϵ* = 10^*−*8^. In the simplified Hebbian t-SNE, the initial values were described as

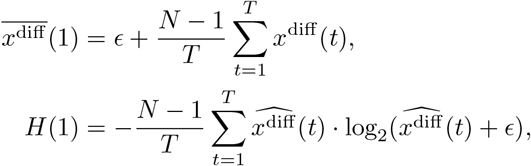

with *ϵ* = 10^*−*8^.

The batch size *T* was set to [*N* (*N −* 1)*/*10], where [x] is a floor function. The number of total steps was 2000 in Figs. 3-5 and Extended Data Fig. 3-6, 10000 in Fig. 2 and Extended Data Fig. 2, 100000 in Fig. 6a-e, 20000 in Fig. 6f-g, and 50000 in Extended Data Fig. 7. The KL divergence plotted in Figs. 2-3 and Extended Data Figs. 2-3 was calculated using the fixed optimal variances *σ*_*j*_ that achieve the target perplexity.

### Original data

The two entangled rings were described as (*r* sin 2*iπ/n, r* cos 2*iπ/n*, 0) and (*r* + *r* sin 2*iπ/n*, 0, *r* cos 2*iπ/n*), where *i* = 0, 1, …, *n−* 1, *r* = 1000 and *n* = 100. The four entangled rings were described as (*r* sin 2*iπ/n, r* cos 2*iπ/n*, 0), 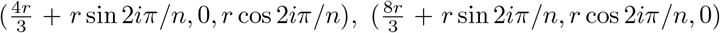, and 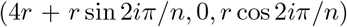, where *i* = 0, 1, …, *n −* 1, *r* = 1000 and *n* = 100. When the reward proportion was *p*, input patterns of *i <* [*np*] were rewarded in Fig. 6a-e.

The S-shaped curve data in Extended Data Fig. 7 included 400 points generated by (150 sin *θ*, 100 sgn(*θ*)(cos *θ−* 1), 200*d*), where sgn(*θ*) represents a sign function, *θ* and *d* were sampled from the uniform distribution between *−*3*π/*2 and 3*π/*2 and the uniform distribution between 0 and 1, respectively. In Fig. 6f-g, input patterns of the 20-40% largest or smallest *θ* values were excluded from the input data. When the reward population was *p*, input patterns of the [200*p*] largest and smallest *θ* values were positive- and negative-rewarded, respectively, in Fig. 6f-g and Extended Data Fig. 7. The MNIST data were constructed by randomly choosing 1200 images from the MNIST dataset.

The experimental data of the receptor responses in Fig. 4 and Extended Data Fig. 4 was obtained from Table S1 in [19]. The experimental data of the PN activities and valence index in Fig. 5 and Extended Data Figs. 5-6 are described in [20]. The raw data of the valence index was provided by the author of [20].

### PCA and t-SNE

The PCA and t-SNE were conducted with the scikit-learn package [65] in Python. The perplexity of the t-SNE was set to 20 in Fig. 2 and 40 in Fig. 3.

### The definition of linear and reward separability

The linear separability was defined as the classification accuracy achieved using a linear support vector machine, implemented by using the scikit-learn package [65]. The reward separability was defined as the maximum accuracy of a classifier that judges the data whose *y*_1_ coordinate exceeds the threshold as belonging to the positive-rewarded group and all other data as belonging to the negative-rewarded group. The threshold was set to maximize the classification accuracy. The positive- and negative-rewarded groups were defined as follows. In Fig. 6a-e, the positive- and negative-rewarded groups encompassed their respective rewarded points as well as unrewarded points within the same rings as the respective rewarded points. In Extended Data Fig. 7, data points were divided into positive- and negative-rewarded groups along the S-shaped manifold as indicated by different colors in Extended Data Fig. 7.

### The analysis related to the odor and valence index

In Fig. 5, we obtained the *K* clusters *C*_1_, *C*_2_, …, *C*_*K*_ by applying the K-means clustering for each representation. We calculated the sum of the squared error *S* between the true valence index and its group mean, described as

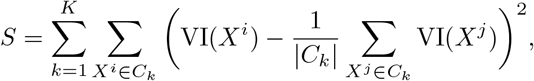

where VI(*X*^*i*^) represents the experimentally measured valence index of the odor *X*^*i*^. We plotted the value of *S* for various numbers of the cluster number *K* in Fig. 5c and Extended Data Fig. 6c. In the shuffled case in Fig. 5c and Extended Data Fig. 6c, the shuffled valence index was used instead of the true valence index.

In Extended Data Figs. 5-6, the normalized Euclidean distances so that the maximum distance was one, were plotted. A one-sided statistical test was used for the Pearson correlation coefficient.

## Acknowledgments

We thank Hokto Kazama for providing the valence index data in [20], valuable comments on the manuscript, and helpful discussions. This study was supported by RIKEN Center for Brain Science (T.T. and K.Y.), JST CREST program JPMJCR23N2 (T.T.), KAKENHI Grant-in-Aid JP21J10564 (K.Y.) and JP23K19415 (K.Y.) from JSPS, and Masason Foundation (K.Y.).

## Author contributions

K.Y. and T.T. conceived the project. K.Y. conducted numerical simulations and data analyses. K.Y. and T.T. wrote the manuscript.

## Data availability

All experimental data used in this study are from the previous studies. The experimental data of the receptor responses in Fig. 4 and Extended Data Fig. 4 are available at [19]. The experimental data of PN activities in Fig. 5 and Extended Data Figs. 5-6 are available at [20]. The experimental data of the valence index in Fig. 5 and Extended Data Figs. 5-6 will be available in a public repository upon publication.

## Code availability

The source code will be available in a public repository upon publication.

## Extended Data for Kensuke Yoshida and Taro Toyoizumi

**Contents**

Extended Data Figs. 1-7.

**Extended Data Fig. 1:**
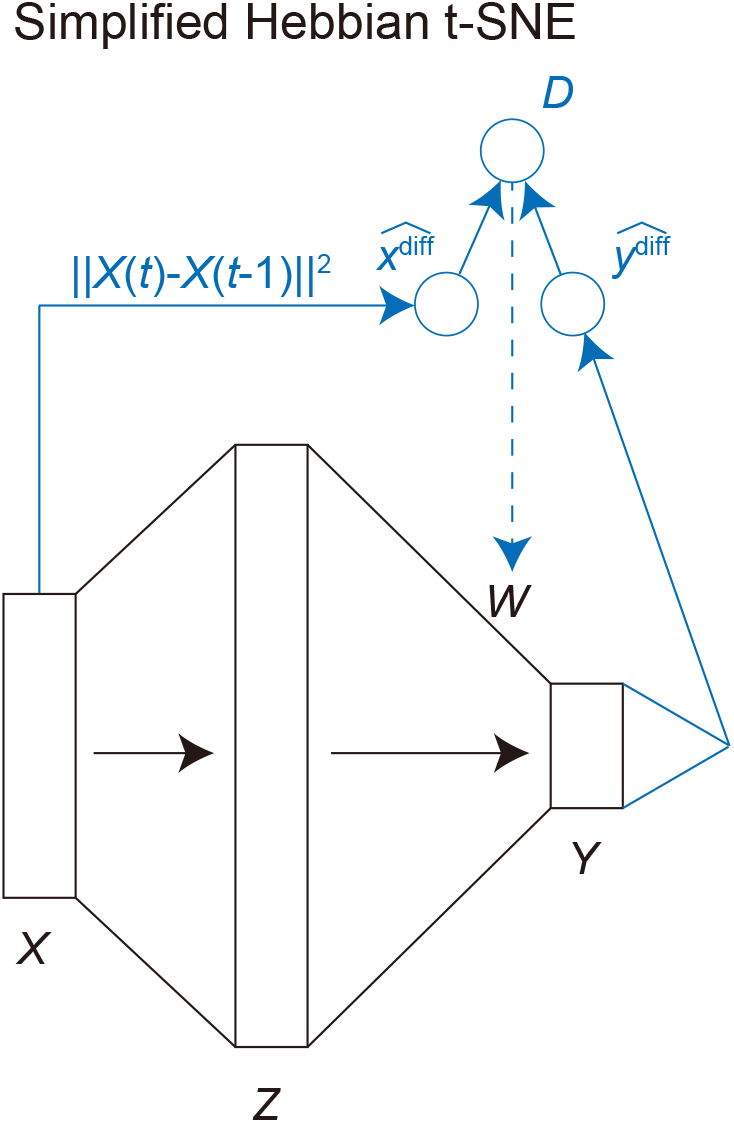
Model structure of the simplified Hebbian t-SNE. The only difference from the original model is that the description of 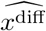 is simplified.

**Extended Data Fig. 2:**
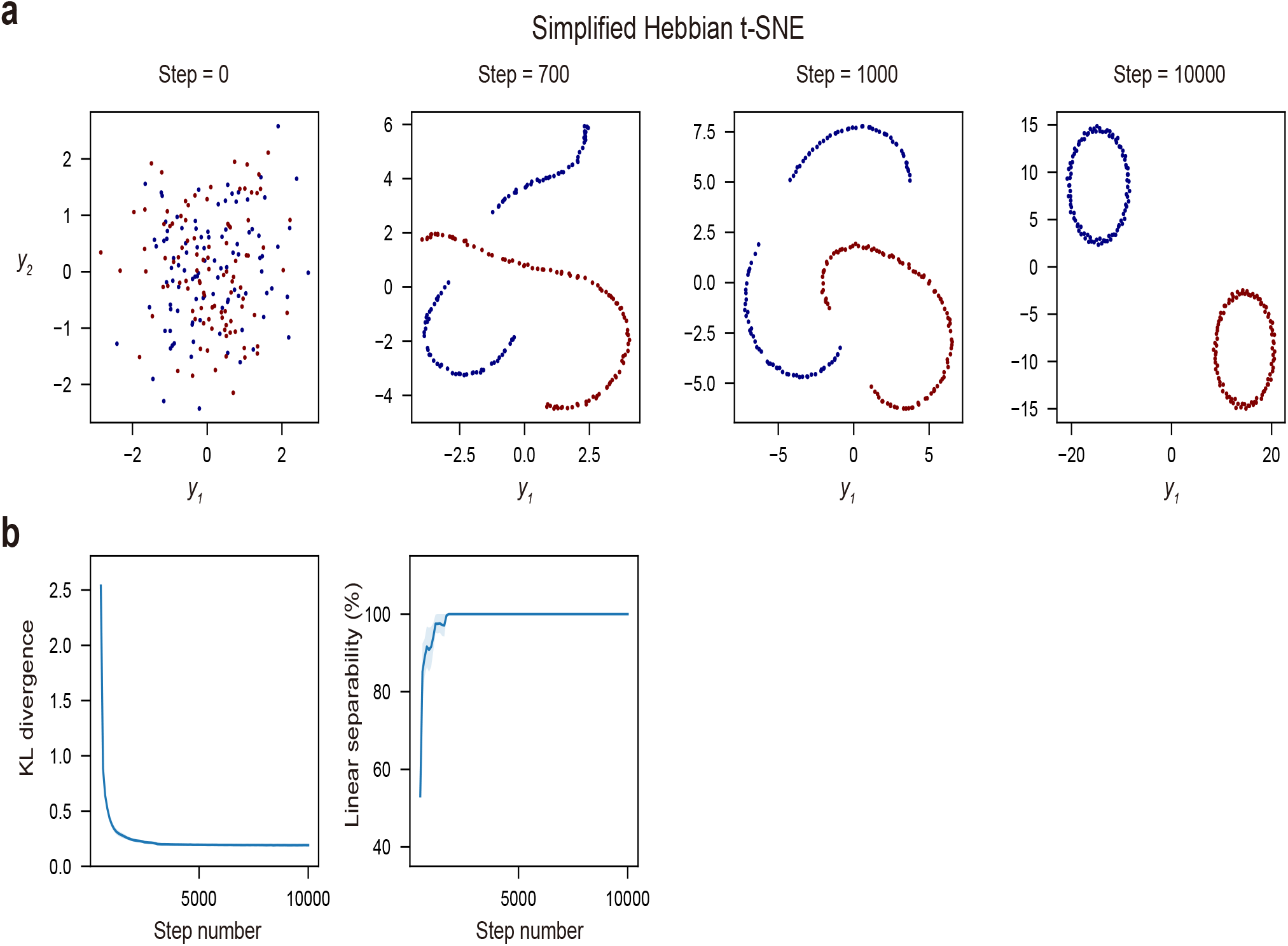
Applying the simplified Hebbian t-SNE to inputs distributed according to entangled rings. (a) The time course of the representation changes by the simplified Hebbian t-SNE. The two rings were gradually separated along the iterative synaptic weight updates as well as the Hebbian t-SNE. (b) The changes of the KL divergence between the input and output similarity matrices and the linear separability of two rings in the output representation. The lines and shadows represent the means and SEMs in the five trials, respectively. **a**

**Extended Data Fig. 3:**
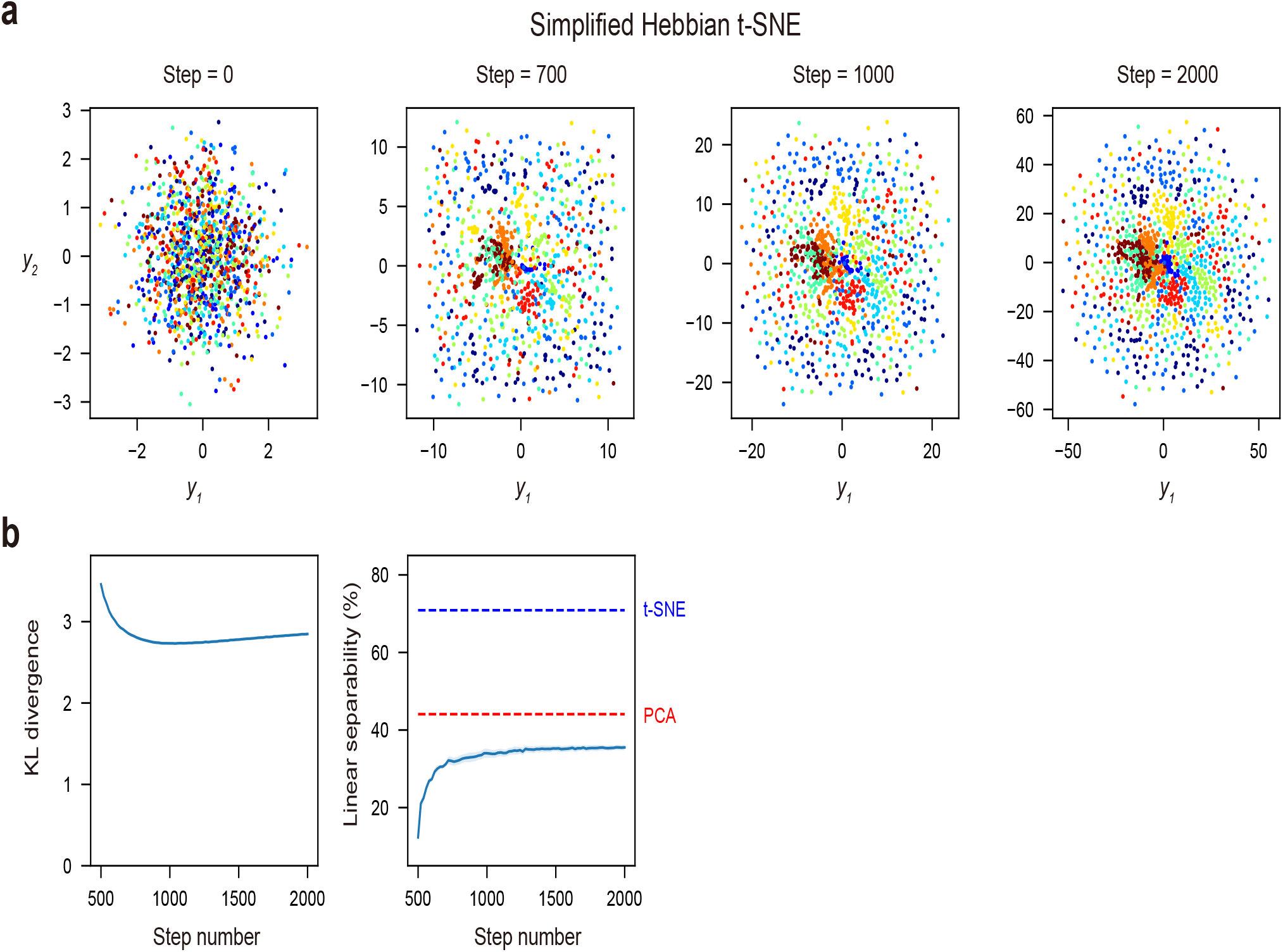
Applying the simplified Hebbian t-SNE to the MNIST data. (a) The time course of the representation changes by the simplified Hebbian t-SNE. The simplified Hebbian t-SNE failed to obtain good representations separating different digits. (b) The changes in the KL divergence between the input and output similarity matrices and the linear separability of the ten digits in the output representation. The lines and shadows represent the means and SEMs in the five trials, respectively. t-SNE

**Extended Data Fig. 4:**
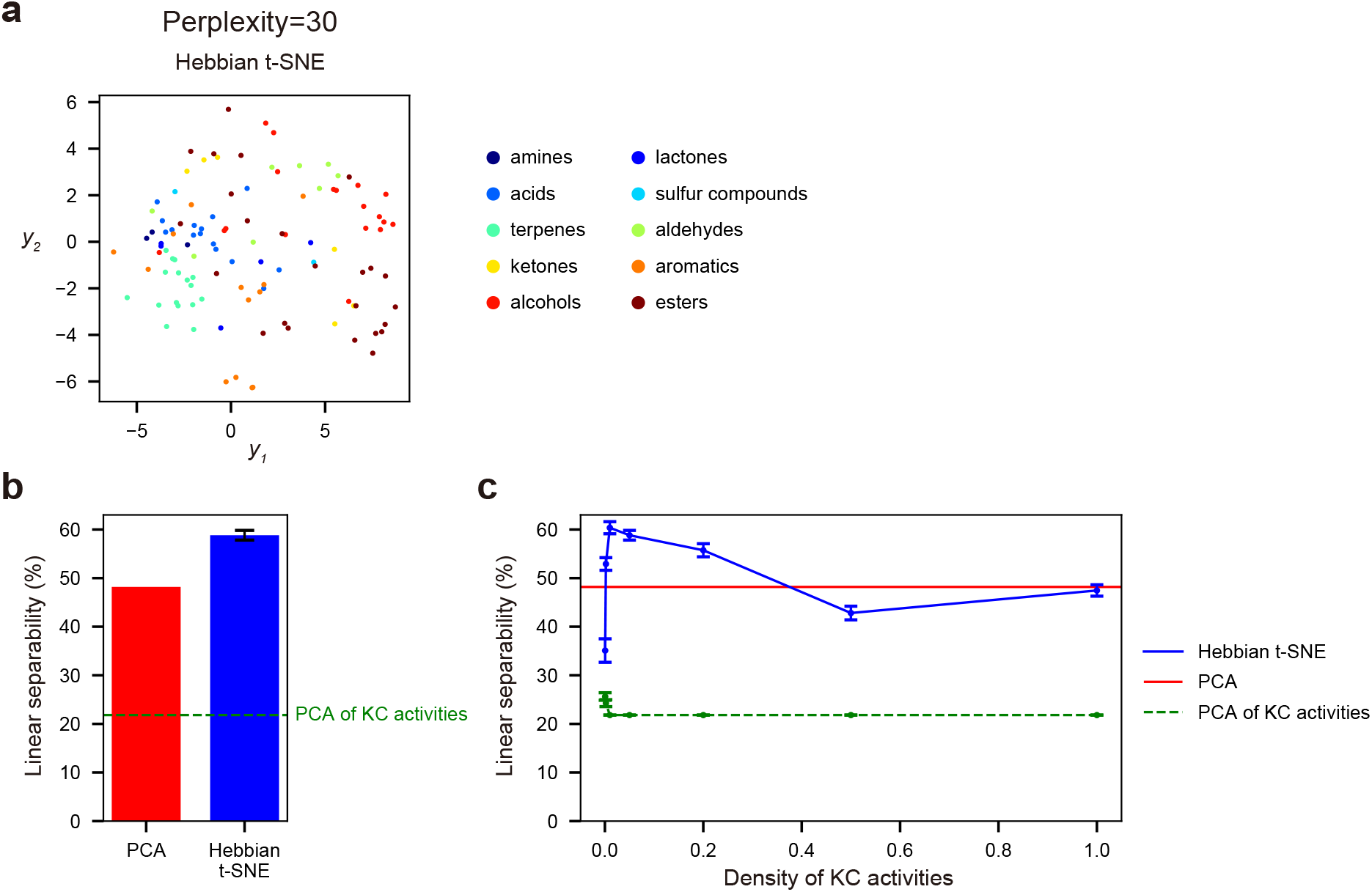
Applying the Hebbian t-SNE in case the perplexity was set to 30, related to Fig. 4. The overall results were the same as Fig. 4. (a) The representations obtained by Hebbian t-SNE. The colors indicate the ten functional groups (amines, lactones, acids, sulfur compounds, terpenes, aldehydes, ketones, aromatics, alcohols, and esters) based on the chemical structures. (b) The linear separability of the ten functional groups in the two-dimensional representations obtained by the PCA and Hebbian t-SNE. The error bar indicates SEMs in the ten trials. The green dotted line indicates the linear separability of the ten functional groups in the two-dimensional representations obtained by applying the PCA to the modeled KC activities. (c) The linear separability of the ten functional groups in the two-dimensional representations when the density of KC activities was varied. The error bars indicate SEMs in the ten trials.

**Extended Data Fig. 5:**
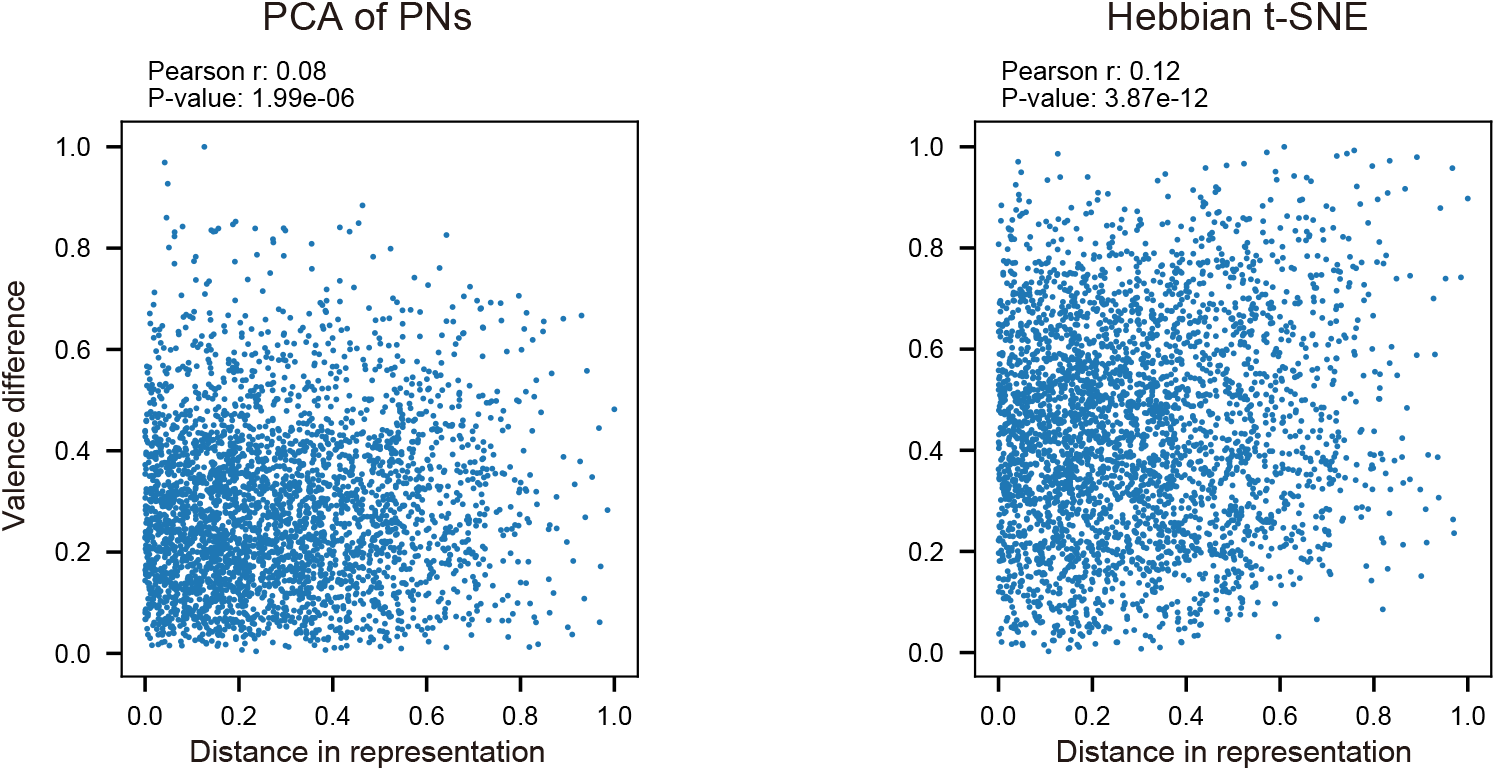
The relationships between the difference in valence index and the Euclidean distance in the two-dimensional representations by the PCA (left) and Hebbian t-SNE (right). The valence difference and Euclidean distance were normalized so that their maximum values were one. A one-sided test was used for the Pearson correlation coefficient.

**Extended Data Fig. 6:**
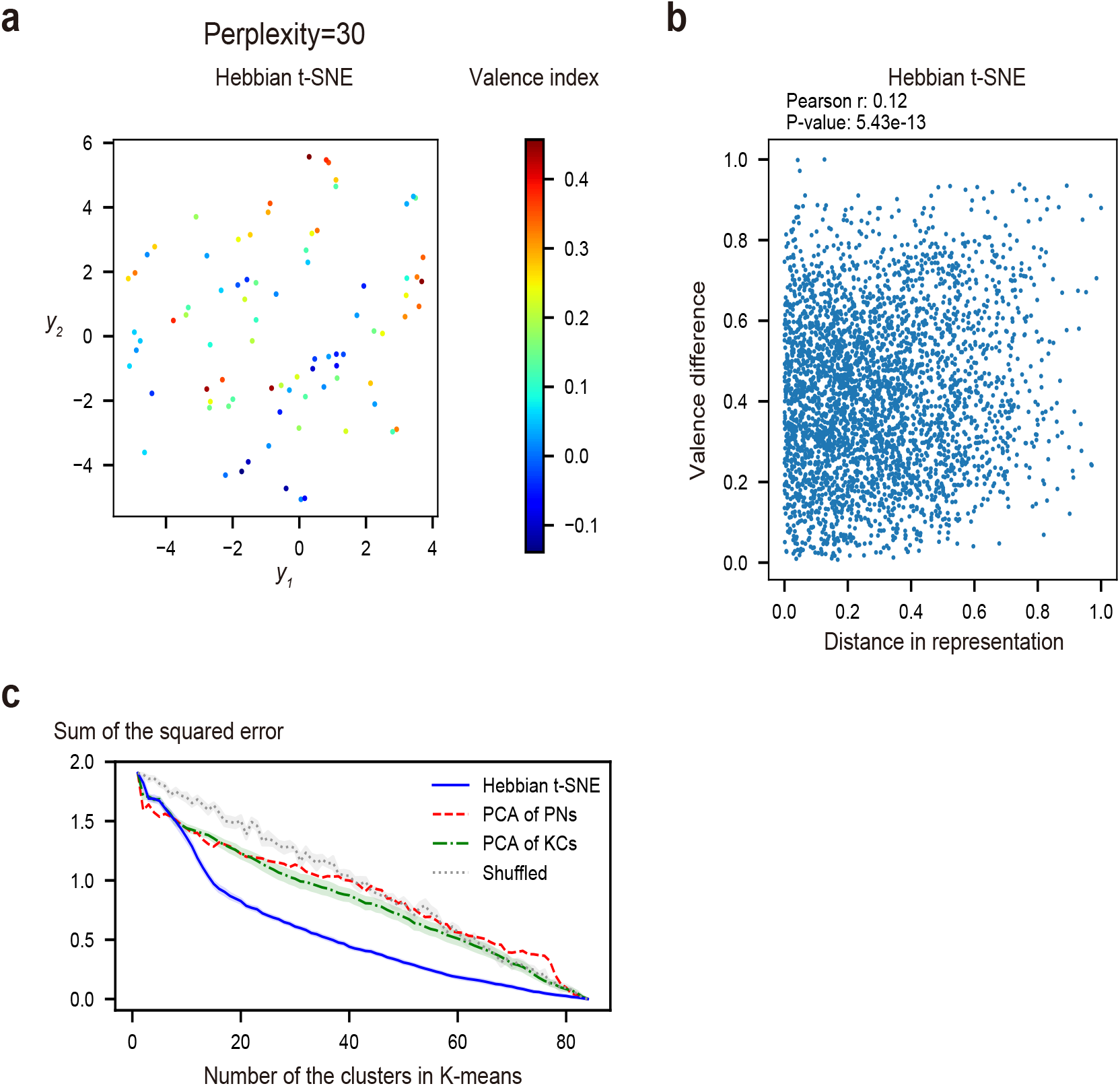
Applying the Hebbian t-SNE in case the perplexity was set to 30, related to Fig. 5 and Extended Data Fig. 5. The overall results were the same as Fig. 5 and Extended Data Fig. 5. (a) The two-dimensional representations by applying the Hebbian t-SNE to the PN activities. The colors indicate the valence index. (b) The relationship between the difference in valence index and the Euclidean distance in the two-dimensional representations by the Hebbian t-SNE. A one-sided test was used for the Pearson correlation coefficient. (c) The squared errors between the valence index of each data point and its neighborhood cluster mean for varying numbers of K-means clusters for each representation. The representations were obtained by applying the PCA to the PN activities (red), PCA to the modeled KC activities (green), and Hebbian t-SNE (blue). The gray line indicates the case of Hebbian t-SNE with the random shuffling of the valence index. The lines and shadows represent the means and SEMs in the ten trials, respectively.

**Extended Data Fig. 7:**
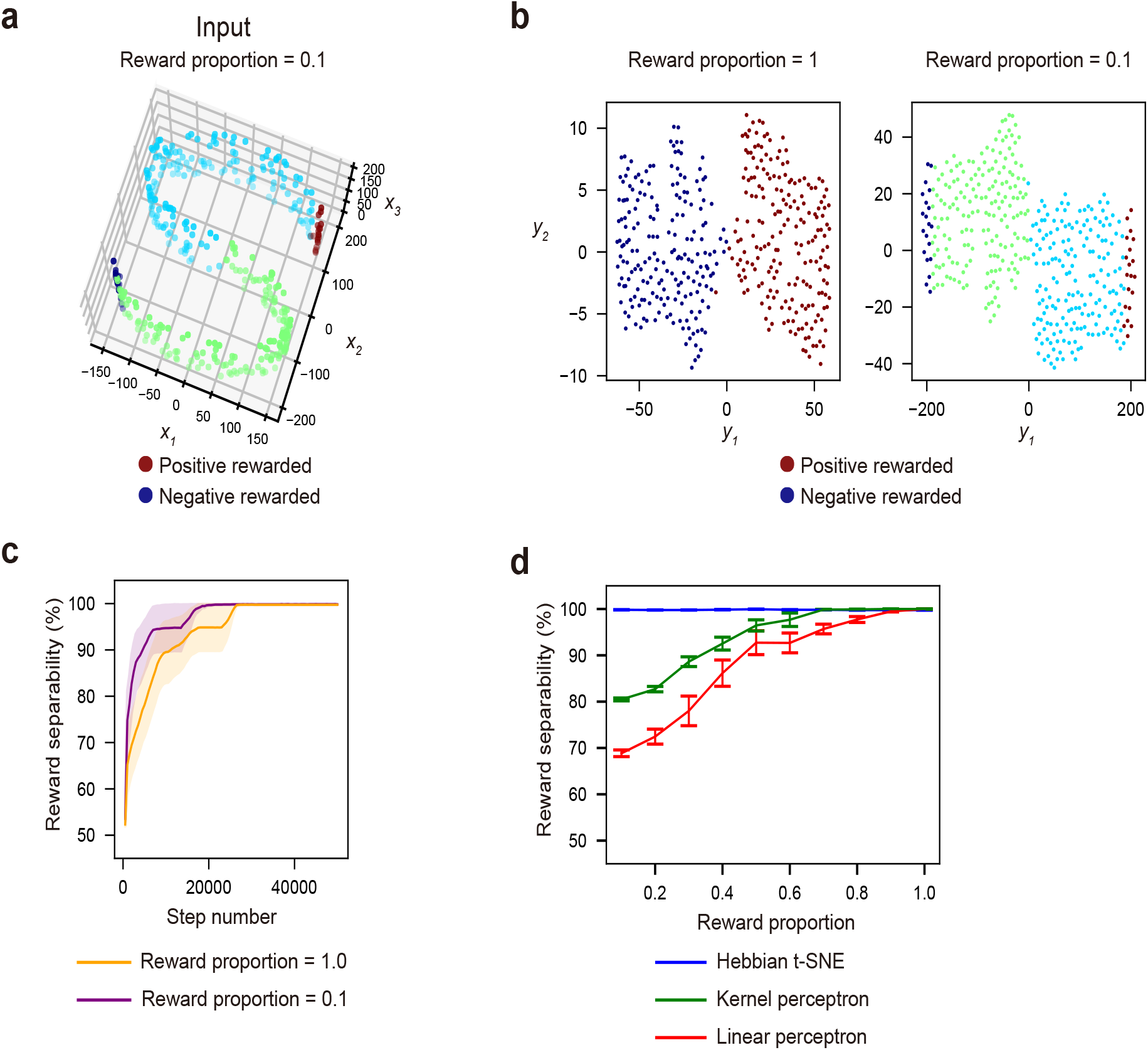
Applying the reward-modulated Hebbian t-SNE to the data in an S-shaped manifold. (a) The original three-dimensional data. The red and blue points indicated positive- and negative-rewarded data, respectively. The reward proportion is the proportion of the rewarded data in the total data. (b) The representations obtained with distinct reward proportions. The red and blue data points accompanied rewards. Their representations were biased to the positive and negative direction of the *y*_1_ coordinate by the positive and negative reward, respectively. (c) The time course of reward separability when the reward proportion is 1.0 (orange) and 0.1 (purple). The lines and shadows represent the means and SEMs in the ten trials, respectively. (d) The reward separability dependent on reward proportions in the Hebbian t-SNE (blue), linear perceptron for three-dimensional inputs (red), and linear perceptron after transforming high-dimensional space (green) (see Methods). The error bars represent SEMs in the ten trials.

